# Nonuniform scaling of cerebellar cortical-nuclear architecture across primates revealed by cross-species atlases

**DOI:** 10.64898/2026.04.01.715860

**Authors:** Kadharbatcha S Saleem, Alexandru V Avram, Daniel Glen, Peter J Basser

## Abstract

The cerebellum shapes distributed motor and association networks through precisely organized pathways linking the cerebellar cortex to the deep cerebellar nuclei (DCN), its principal output structures. Whether these cortical and nuclear compartments scale proportionally across primates and how their relative expansion influences large-scale brain organization remains unclear. We generated multimodal, high-resolution cross-species 3D cerebellar atlases in marmoset, macaque, and human, integrating iron-sensitive magnetic resonance contrast with complementary histological markers to delineate DCN subdivisions in detail. Comparative volumetric analyses reveal nonuniform scaling of cerebellar cortical-nuclear architecture: cortical expansion markedly outpaces enlargement of the DCN, indicating disproportionate growth of input relative to output structures. This divergence parallels preferential expansion of posterior hemispheric territories, particularly lobules VI-IX and Crus I/II linked to higher-order association networks, whereas DCN subdivisions show selective rather than uniform scaling. Together, these findings establish nonuniform cortical-nuclear scaling as a systems-level organizational principle that reshapes cerebellar contributions to distributed brain networks.

## Introduction

The cerebellum is increasingly recognized as a central node in distributed brain systems supporting sensorimotor, cognitive, and affective functions (*1–4*). Once considered primarily as a motor structure, converging anatomical, physiological, and neuroimaging evidence demonstrates that cerebellar output modulates widespread cortical and subcortical networks through precisely organized cortico-nuclear-thalamic pathways (*5, 6*). Within this architecture, the cerebellar cortex performs extensive internal computations, whereas the deep cerebellar nuclei (DCN) constitute the sole efferent channels transmitting cerebellar signals to motor, premotor, associative, and limbic circuits (*7, 8*). The relative scaling between the cerebellar cortical sheet and its nuclear output structures, therefore, imposes a structural constraint on how cerebellar computations are distributed across large-scale brain systems.

Comparative studies in primates document marked cerebellar enlargement, particularly within hemispheric territories functionally coupled to higher-order cortical regions (*9–11*). Posterior lobules, including lobules VI-IX and the ansiform lobule (Crus I/II), are disproportionately expanded in anthropoid primates and are consistently engaged during human cognitive tasks (*1, 12*). These findings implicate posterior cerebellar territories in large-scale association networks and support a cerebellar role extending beyond classical motor control. However, most comparative analyses emphasize gross cerebellar size or surface expansion, leaving unresolved whether expansion of the cerebellar cortical sheet is accompanied by proportional scaling of the DCN. Because the DCN constitute the exclusive efferent interface of the cerebellum, disproportionate scaling between cortical territories and their nuclear output compartments would imply structural reconfiguration of cerebellar influence across distributed systems.

The DCN are structurally and functionally differentiated into dentate, interposed, and fastigial nuclei, each defined by distinct connectivity and computational roles (*7, 8*). In primates, the dentate nucleus exhibits pronounced morphological elaboration consistent with expanded functional domains beyond primary motor output (*13, 14*). Yet detailed cross-species characterization of DCN subdivisions and their relative scaling remains limited, in part because conventional imaging approaches do not fully resolve fine nuclear boundaries. Iron-sensitive magnetic resonance imaging (MRI) provides contrast sensitive to cytoarchitectural and myeloarchitectural variation within the DCN (*15–17*), raising the possibility that species-dependent iron distribution patterns reflect systematic differences in nuclear organization.

Resolving these questions requires multimodal cerebellar atlases that preserve anatomical fidelity while enabling quantitative comparison of cerebellar cortical and nuclear domains. Integration of high-resolution MRI with histological validation permits anatomically precise region mapping suitable for cross-species alignment (*18–20*). Establishing such atlases across primates provides a foundation for determining how cerebellar input and output compartments scale within hierarchical brain systems.

To address this, we generated multimodal, high-resolution cross-species 3D cerebellar atlases in marmoset, macaque, and human. Using atlases combining iron-sensitive and immunohistochemical contrasts, we delineate DCN subdivisions and characterize species-dependent structural differentiation of cerebellar output systems. We quantify volumetric variation across DCN subdivisions to determine whether cerebellar output structures scale uniformly or exhibit selective expansion. Finally, we examine whole-cerebellar and lobular volumes to assess cortical scaling relative to nuclear output structures, with particular emphasis on posterior hemispheric territories. This hierarchical analysis from nuclear microstructure to global cerebellar organization reveals nonuniform scaling of cerebellar output structures alongside progressive posterior hemispheric specialization, establishing a systems-level framework for understanding how cerebellar architecture supports distributed motor and association networks across primates.

## Results

Multimodal MRI and histology were integrated to generate high-resolution cross-species cerebellar atlases in marmoset, macaque, and human, enabling quantitative comparison of lobular territories and deep cerebellar nuclei (DCN). Iron-sensitive and immunohistochemical contrasts delineated DCN subdivisions across species and revealed distinct patterns of nuclear differentiation. Volumetric analyses demonstrated that cerebellar cortical expansion markedly exceeded DCN enlargement, indicating disproportionate scaling of input relative to output compartments. This dissociation was accompanied by progressive enlargement of posterior hemispheric territories, particularly lobules VI-IX and Crus I/II, whereas nuclear subdivisions exhibited selective rather than global expansion. Together, these findings demonstrate nonuniform scaling of cerebellar input-output architecture across primates.

### Distinct patterns of cerebellar iron accumulation across primates

Using high-resolution MRI validated with histology (see Methods; Fig. 9), we examined interspecies differences in cerebellar signal and iron distribution, focusing on the deep cerebellar nuclei (DCN). Pronounced differences in MR signal were observed across species, especially within the DCN. On T2-weighted images, the DCN appeared strongly hypointense relative to surrounding cerebellar tissue in macaques and humans, whereas this contrast was markedly reduced or hyperintense in marmosets (Fig. 1A, B, E). These differences are consistent with prior studies linking T2 hypointensity to elevated iron content in neural tissue (*21, 22*). Histological analysis using Perls’ Prussian blue staining confirmed these patterns: macaque DCN showed robust iron labeling throughout the neuropil, whereas marmoset DCN displayed minimal staining (Fig. 1C, D), indicative of lower iron levels. The close correspondence between MRI contrast and histologically measured iron demonstrates a species-dependent relationship between DCN iron accumulation and T2-weighted signal properties. These results reveal distinct patterns of cerebellar iron distribution across primates and provide a mechanistic framework for interpreting cross-species differences in MR signal, with implications for translational neuroimaging.

**Fig. 1.**
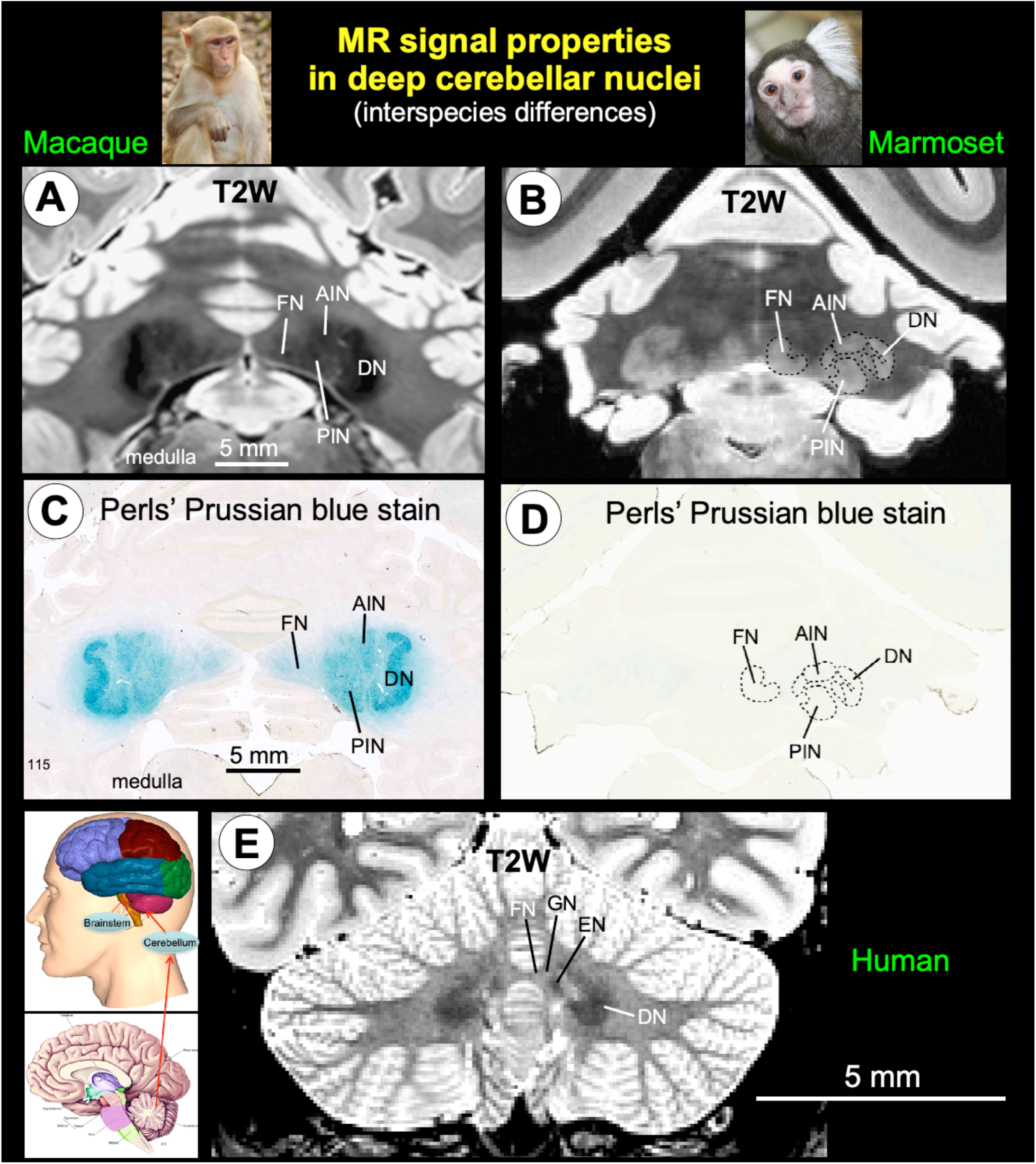
Comparative anatomy and iron distribution of the deep cerebellar nuclei across primates. **A-B**: T2-weighted MRI of the deep cerebellar nuclei (DCN) in macaque and marmoset, showing the dentate nucleus (DN), anterior interposed nucleus (AIN or EN-Emboliform nucleus), posterior interposed nucleus (PIN or GN-globose nucleus), and fastigial nucleus (FN). **C-D**: Corresponding histological sections stained with Perls’ Prussian blue to detect ferric iron. In macaques, DCN display strong iron labeling, consistent with the hypointense T2 signal, whereas marmoset DCN show minimal iron staining and relative hyperintensity on MRI. **E**: T2-weighted image of the human DCN (from in vivo, Connectome 1.0) exhibits hypointense signals similar to macaque, reflecting higher iron content. These data illustrate species-specific differences in DCN iron accumulation and demonstrate correspondence between MRI contrast and histological iron distribution.

### Histology-validated mapping of deep cerebellar nuclei across non-human primates

The deep cerebellar nuclei (DCN): dentate (DN), anterior interposed (AIN; or emboliform nucleus, EN), posterior interposed (PIN; or globose nucleus, GN), and fastigial (FN) were delineated in marmosets and macaques using multimodal MRI (T2-weighted, MTR, and MAP-MRI) with matched histological validation (SMI-32, NeuN, AChE, and Nissl; Figs. 2, 3). At 150-200 μm isotropic resolution, T2-weighted imaging reliably resolved boundaries among the DN, interposed nuclei, and FN (Fig. 2A, B; figs. S1 and S2), capturing key features of DCN compartmental organization. In the marmoset, the DN appeared partially continuous with the AIN at select rostrocaudal levels (fig. S1, section 96). In this species, MTR and MAP-MRI metrics (radial diffusivity [RD] and return-to-axis probability [RTAP]) provided clear contrast among DCN subregions. In the macaque, MTR maintained discernible nuclear contrast, whereas RD and RTAP showed limited differentiation (Fig. 3).

**Fig. 2.**
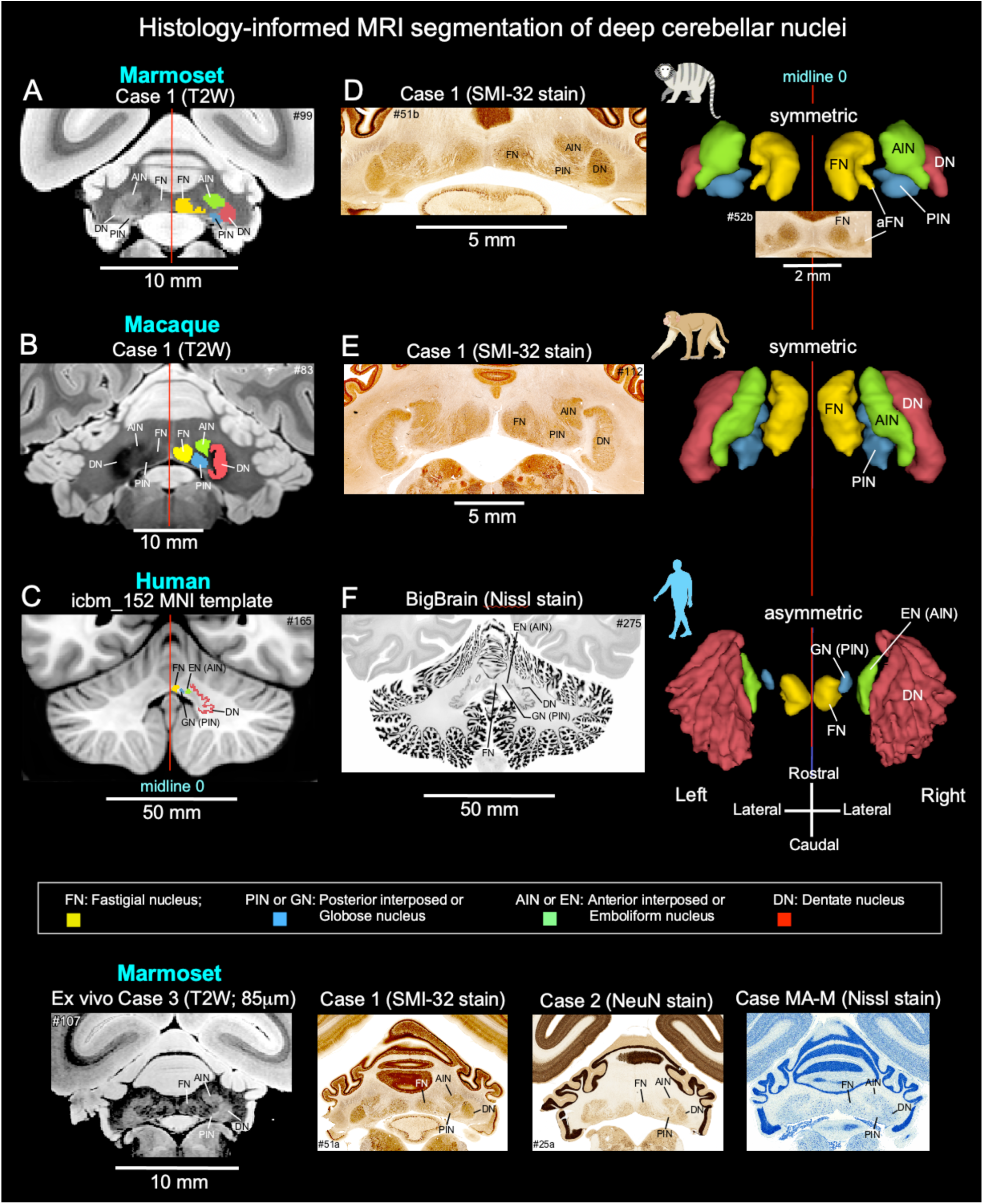
Histology-guided segmentation of the deep cerebellar nuclei (DCN) across primate species. **A-C**: Delineation of the dentate nucleus (DN), anterior interposed nucleus (AIN; or emboliform nucleus, EN), posterior interposed nucleus (PIN; or globose nucleus, GN), and fastigial nucleus (FN) in high-resolution T2-weighted (T2W) MRI of marmoset (150 μm; case 1) and macaque (200 μm; case 1), and in the human population-based MNI template (500 μm). **D-E**: MRI-defined DCN borders were validated against matched histological sections from the same specimens stained with SMI-32, demonstrating concordant nuclear boundaries and internal architecture. (**F**) For humans, the BigBrain Project dataset was registered to the MNI template to enable histology-informed delineation of DCN subregions, including the highly folded and crenated DN, which is not reliably resolved on T1-weighted MNI templates alone. **Right column**: Three-dimensional reconstructions illustrating the rostrocaudal extent of the DCN across species (superior view). In the marmoset, a small satellite structure contiguous with the FN was consistently identified and designated the accessory fastigial nucleus (aFN) (top right; inset shows magnified SMI-32-stained section). **Bottom row**: DCN delineation in an independent high-resolution (85 μm) T2W dataset (marmoset case 3) with closely matched histological sections from independent cases stained with SMI-32, NeuN, and Nissl. A distinct cell-sparse zone separating the DN from the AIN is clearly resolved at this resolution, in contrast to case 1 (**A**). Across species and imaging modalities, DCN subregions exhibited close correspondence in shape, spatial location, and internal cytoarchitecture between MRI and histology.

**Fig. 3.**
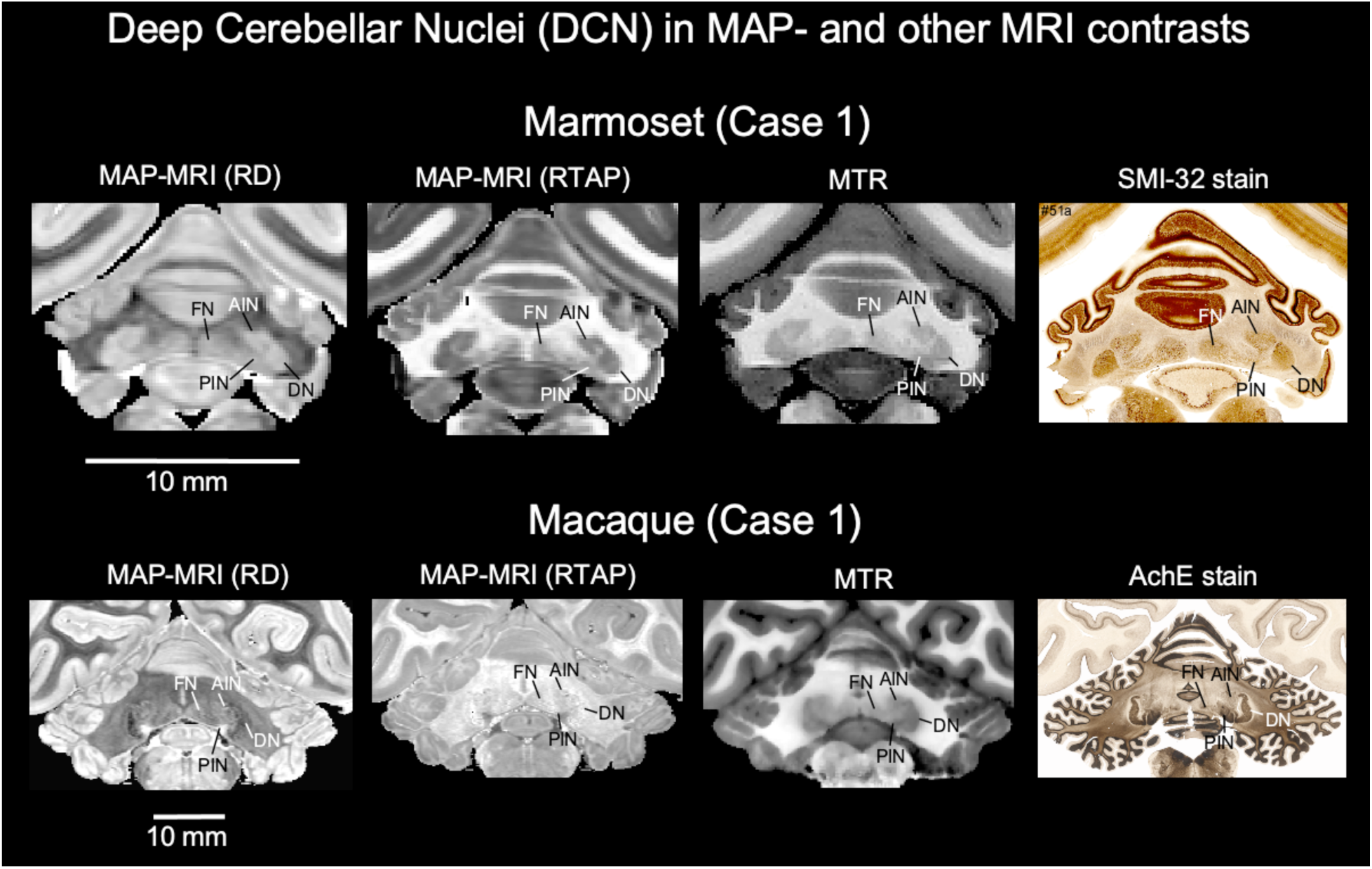
Deep cerebellar nuclei (DCN) across MRI contrasts. Visualization of the dentate nucleus (DN), anterior interposed nucleus (AIN; or emboliform nucleus, EN), posterior interposed nucleus (PIN; or globose nucleus, GN), and fastigial nucleus (FN) using MAP-MRI-derived metricsradial diffusivity (RD) and return-to-axis probability (RTAP), and additional contrasts (magnetization transfer ratio [MTR]) in marmoset and macaque. Matched histological sections from the same specimens are shown for reference (SMI-32 in marmoset; acetylcholinesterase [AChE] in macaque). In the marmoset, RD, RTAP, and MTR provided a clear contrast for distinguishing DCN subregions. In the macaque, MTR maintained discernible nuclear contrast, whereas RD and RTAP showed only limited differentiation between nuclei.

Histology confirmed and refined MRI-based observations. SMI-32 immunostaining distinguished the DN from the interposed nuclei in both species (Fig. 2D, E; figs. S1 and S2). In marmosets, stronger SMI-32 reactivity within the DN neuropil, together with a cell-sparse zone separating it from the AIN and PIN, facilitated clear boundary identification. In macaques, a prominent cell-sparse zone between the DN and interposed nuclei enabled reliable separation despite more uniform labeling intensity. These patterns indicate that nuclear segregation is supported by distinct microanatomical features across species. Notably, in marmosets, a small satellite structure contiguous with the FN was consistently observed and designated the accessory fastigial nucleus (aFN) (Fig. 2, top right; inset). Three-dimensional MRI-histology reconstructions illustrate the rostrocaudal extent and spatial relationships of DCN subregions across species, highlighting differences in relative expansion and folding (Fig. 2, right column). In one marmoset (case 3), higher-resolution T2-weighted imaging (85 μm isotropic) revealed a narrow cell-sparse zone separating the DN from the AIN and adjacent nuclei. Although direct histological validation was unavailable for this specimen, MRI-defined boundaries corresponded closely with patterns observed in SMI-32, NeuN, and Nissl-stained sections from independent animals (Fig. 2, bottom row).

### Mapping human deep cerebellar nuclei in a comparative primate framework

In humans, DCN subregions were delineated in MNI space (500 μm resolution) using the BigBrain Project dataset, derived from ultrahigh-resolution Nissl-stained histological sections registered to the MNI template (Fig. 2C, F; fig. S3). Conventional T1-weighted MNI templates alone did not reliably resolve DCN boundaries. BigBrain-informed segmentation enabled clear identification of all major DCN subregions, including the highly folded dentate nucleus, while preserving the rostrocaudal ordering of nuclei relative to cerebellar lobules. To enable quantitative analyses beyond the MNI reference, four additional in vivo T2-weighted datasets (800 μm resolution) were analyzed. In these cases, DCN were delineated rostrocaudally with reference to corresponding BigBrain histology registered to each T2 volume, extending the histology-based framework to living human brains and providing independent datasets for volume and morphometric measurements (Fig. 4, top; fig. S4).

**Figure 4.**
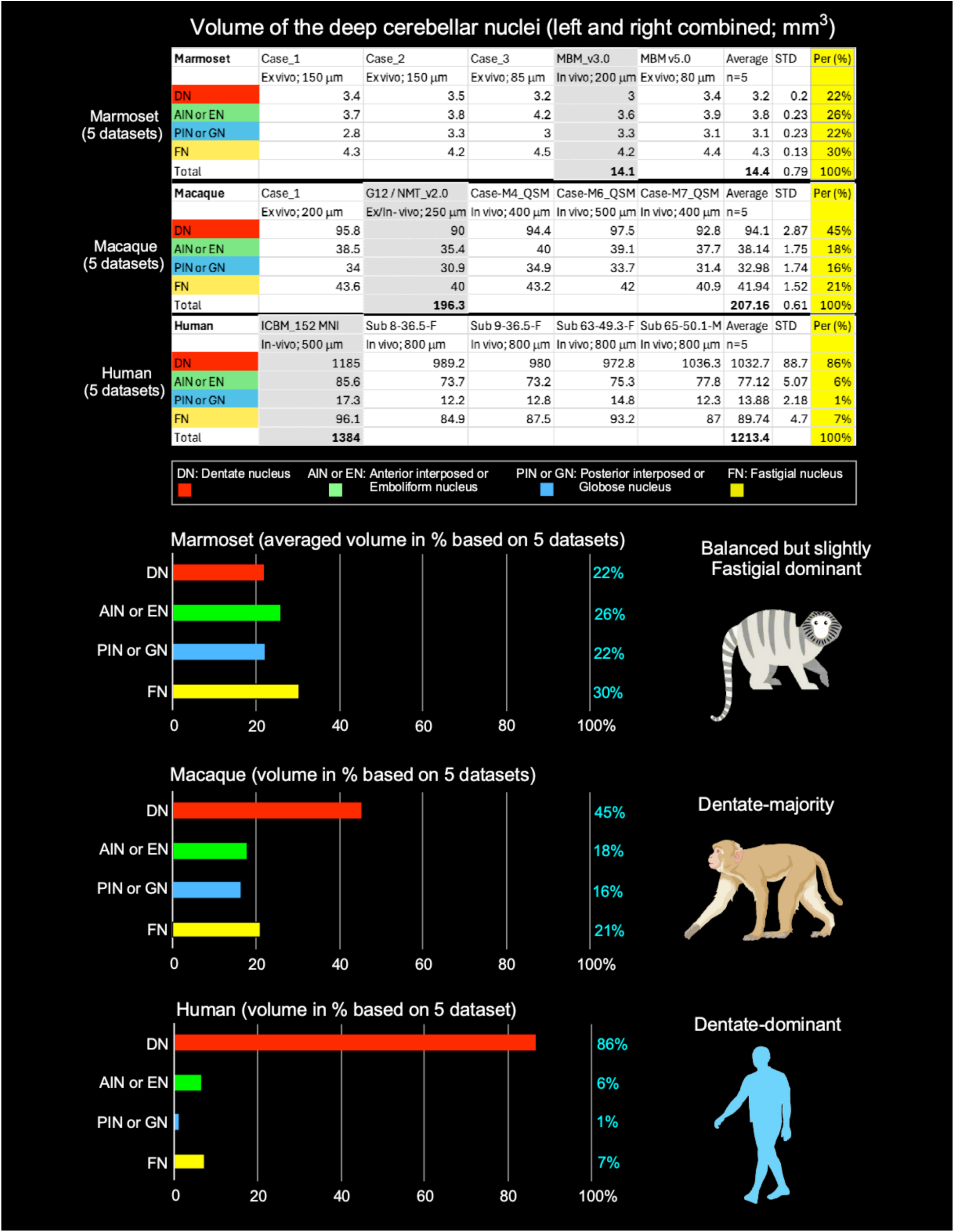
Volume of the deep cerebellar nuclei (DCN). **Top:** Absolute volumes of DCN subregions (left and right combined) in marmosets, macaques, and humans, averaged across five individuals per species. Individual case information, including species, age, and MRI acquisition parameters (ex vivo or in vivo) with spatial resolution, is summarized in the accompanying figure and detailed in Materials and Methods. For human datasets, one case was acquired at 500 μm isotropic resolution, whereas the remaining four cases were acquired at 800 μm isotropic resolution. The BigBrain (histology) atlas was registered to each subject to guide segmentation of the DCN. **Bottom:** For cross-species comparison, subregion volumes are expressed as a percentage of total DCN volume.

### Scaling of deep cerebellar nuclei across primates

We quantified the relative volumes of deep cerebellar nuclei (DCN) subregions in marmosets, macaques, and humans, averaging measurements from five individuals per species (Fig. 4, top). To enable direct cross-species comparison despite marked differences in absolute brain size, volumes are expressed as percentages of total DCN volume (Fig. 4, bottom).

In marmosets, DCN subregions were relatively evenly distributed. The fastigial nucleus (FN) represented the largest fraction of the DCN (30%), followed by the anterior interposed/emboliform nucleus (AIN/EN; 26%). The dentate nucleus (DN) and posterior interposed/globose nucleus (PIN/GN) contributed similar proportions (∼22% each). In macaques, the DN showed a notable relative expansion, accounting for 45% of total DCN volume, while FN contributed 21%, and the AIN/EN and PIN/GN comprised 18% and 16%, respectively. Compared with marmosets, macaques displayed a marked increase in the proportional size of the DN and a corresponding reduction of the interposed nuclei.

In humans, the DCN were overwhelmingly dominated by the dentate nucleus (DN, 86% of total DCN), with the AIN/EN and FN contributing 6% and 7%, respectively, and the PIN/GN comprising only 1% (Fig. 4). Notably, DN volume in the ICBM152 MNI template (1,185 mm³) exceeded that in four individual T2-weighted human cases (972.8-1,036.3 mm³; mean 994.6 mm³), representing an approximate 19% increase likely due to template averaging effects that exaggerate small, high-contrast structures. Together, these findings reveal a progressive, species-dependent reorganization of DCN composition, characterized by disproportionate expansion of the DN across primates, with the most pronounced enlargement observed in humans.

Cross-species comparisons reveal a systematic evolutionary trend: DN relative volume increases from 22% in marmosets to 45% in macaques and 86% in humans, whereas the proportional contributions of the interposed nuclei (AIN/EN and PIN/GN) and the FN decline, with the PIN/GN showing the most pronounced reduction (22% → 16% → 1%). These findings highlight a striking, evolutionarily conserved reorganization of DCN subregions, emphasizing the human-specific expansion of the dentate nucleus, consistent with its enhanced role in cerebellar output to higher-order cortical networks.

### Species-dependent scaling of deep cerebellar nuclei relative to total cerebellar size

For total DCN volume, we used values derived from in vivo population-based datasets in each species (Marmoset: 14.1 mm³; Macaque: 196.3 mm³; Human: 1,384 mm³; Fig. 4, top panel; gray-highlighted) to ensure consistency with total cerebellar volume estimates (Table 2). This corresponds to an approximate 13.9-fold increase from marmoset to macaque and a further ∼7.0-fold increase from macaque to human, yielding an overall ∼98-fold expansion. Individual DCN subcomponents were quantified as averaged volumes across specimens. Together, these results indicate a nonuniform reorganization of cerebellar output structures across primates, characterized by disproportionate expansion of specific subcomponents, particularly the dentate nucleus in humans.

When normalized to total cerebellar volumes derived from the same population-based datasets (Table 2; Marmoset: 599.14 mm³; Macaque: 6,685.42 mm³; Human: 154,161.1 mm³), distinct species-dependent scaling patterns emerge (Fig. 5). DCN comprise ∼2.35% of total cerebellar volume in marmosets, ∼2.94% in macaques, and ∼0.90% in humans, indicating that, despite substantial absolute expansion, DCN volume does not scale proportionally with cerebellar size in humans. These findings further support a nonuniform scaling of cerebellar output structures, consistent with the pronounced dominance of the dentate nucleus in humans.

**Figure 5.**
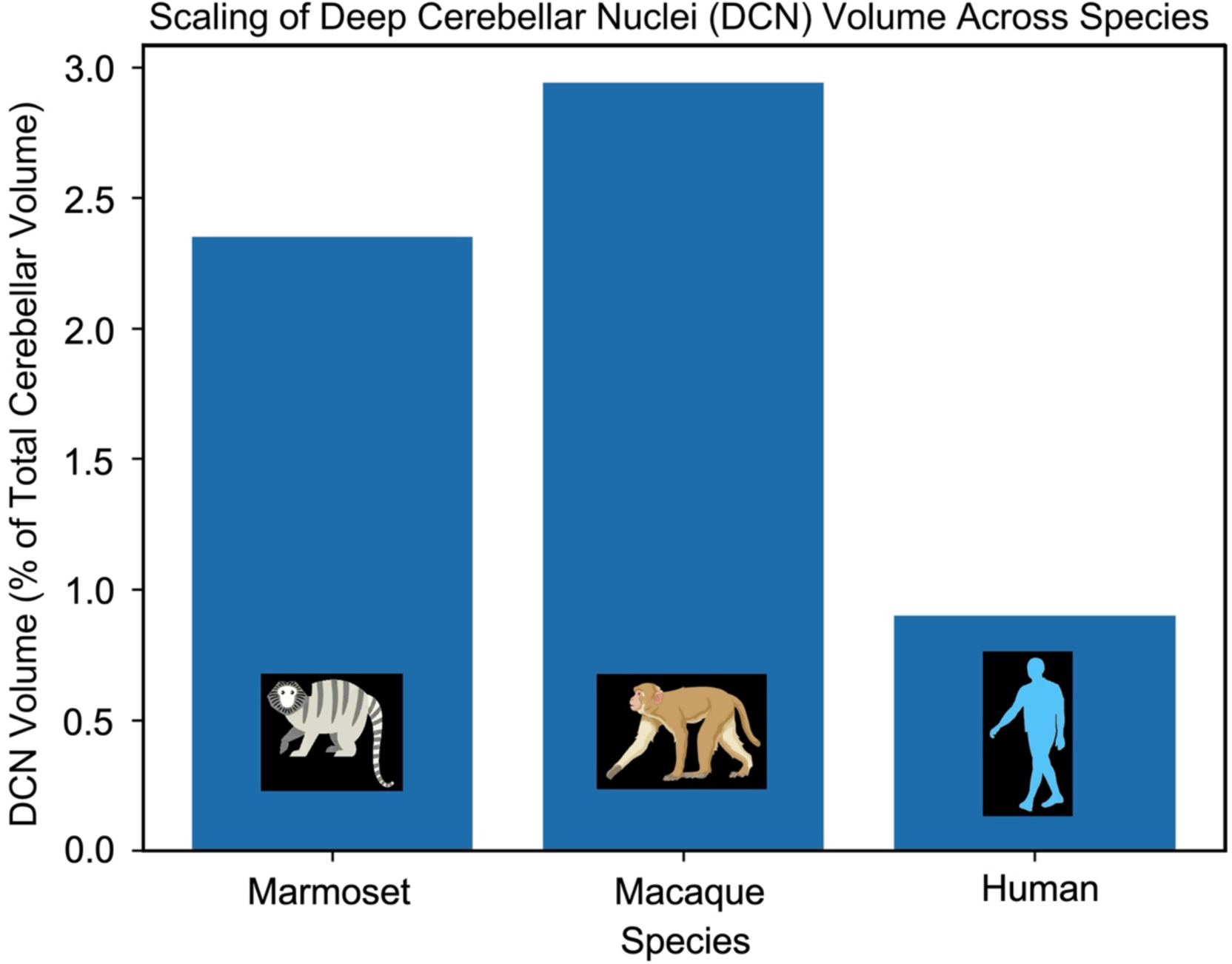
Scaling of DCN relative to total cerebellar volume across primates. DCN volume expressed as a fraction of total cerebellar volume reveals nonproportional scaling across species. DCN comprises ∼2.35% of the cerebellum in marmosets and ∼2.94% in macaques, but only ∼0.90% in humans. This pattern shows that, although DCN volume increases in absolute terms, it represents a smaller proportion of total cerebellar volume in humans compared to nonhuman primates.

### Cerebellar Lobular Morphology and Volume Across Primates

#### Cross-species cerebellar atlases reveal progressive hemispheric differentiation

The disproportionate expansion of cerebellar cortex relative to the deep cerebellar nuclei across primates prompted us to examine whether this differential scaling is accompanied by systematic reorganization of posterior hemispheric architecture. To address this, we leveraged multimodal three-dimensional cerebellar atlases for marmoset, macaque, and human generated from combined high-resolution MRI and histology (see Materials and Methods). These atlases provide precise volumetric delineation of vermal and hemispheric lobules, as well as the deep cerebellar nuclei, using standardized Larsell and Schmahmann nomenclature (*23–25*) and visualization across coronal, axial, and sagittal planes in stereotaxic and MNI space (Figs. 6, 7). The human cerebellar atlas (we call it “HCA”) further incorporates detailed fissural and lobular definitions (*25*) and also provides a flat map of the cerebellum derived from the SUIT cerebellum (*26*) with asymmetric parcellation from HCA (Fig. 7). Together, these resources enable systematic cross-species comparison of lobular boundaries while maintaining species-specific morphology.

**Figure 6.**
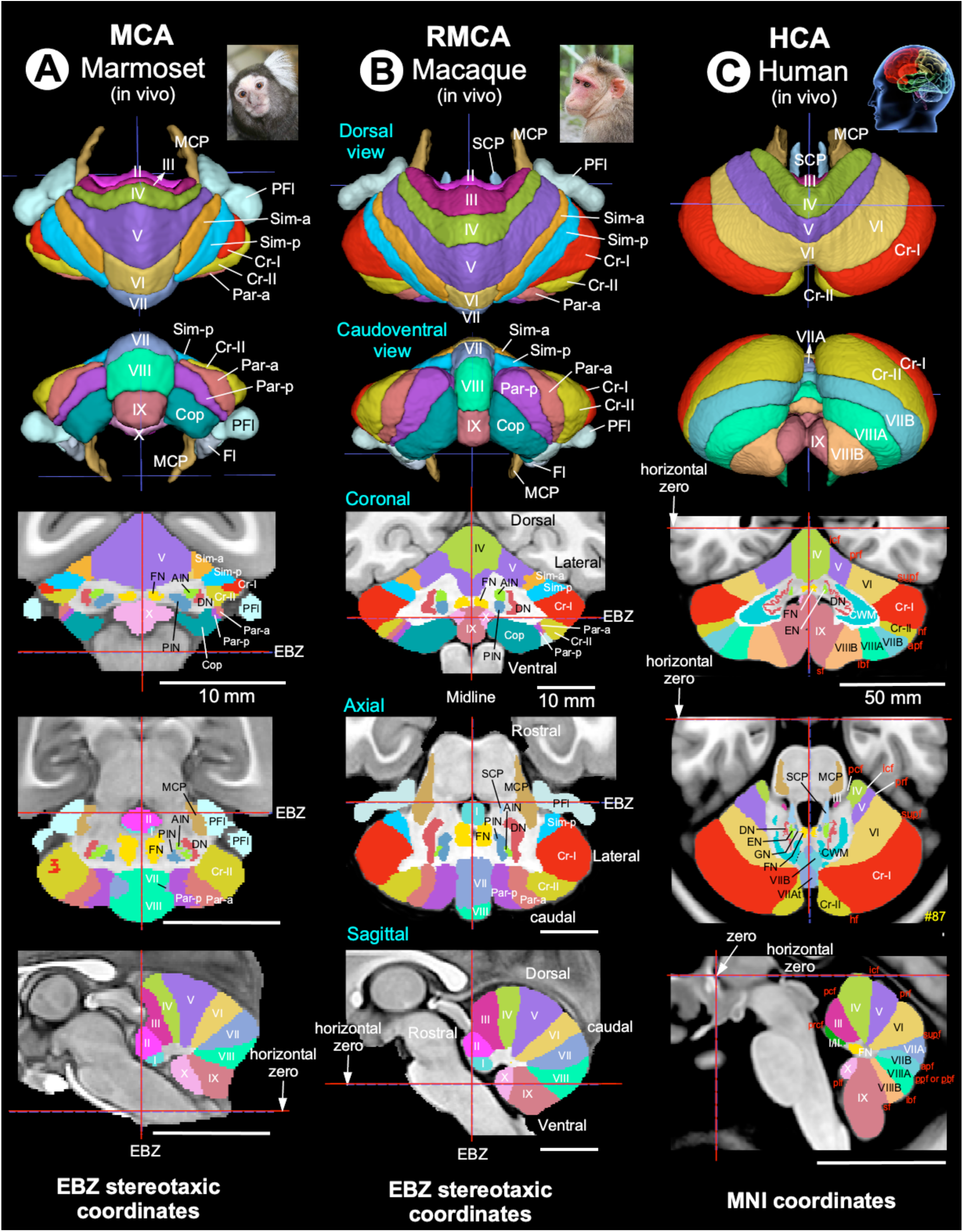
Cross-species 3D cerebellar atlases. High-resolution, multimodal cerebellar atlases were generated for marmoset, macaque, and human by integrating MRI and histology (see Materials and Methods). For marmoset and macaque, ex vivo segmentations of cerebellar lobules, deep cerebellar nuclei, and cerebellar peduncles were transformed into species-specific stereotaxic space (Ear Bar Zero: EBZ) and aligned to population-based in vivo templates to produce the final atlases. These reconstructions form the basis of the Marmoset Cerebellar Atlas (MCA) and Rhesus Macaque Cerebellar Atlas (RMCA) and serve as standard templates for quantitative analyses (**A, B**). For humans, cerebellar segmentation was performed on the in vivo ICBM_152 MNI template with reference to the histological BigBrain dataset registered to MNI space, forming the Human Cerebellar Atlas (HCA) (**C**). These atlases enable precise volumetric delineation of vermal and hemispheric lobules and the deep cerebellar nuclei, standardized using Larsell and Schmahmann nomenclature (*23–25*). Visualization across coronal, axial, and sagittal planes in stereotaxic and MNI spaces facilitates direct cross-species comparisons of cerebellar architecture. ***Abbreviations:*** I-X: cerebellar lobules; AIN: anterior interposed nucleus; Cop: copula of pyramis; Cr-1: crus I (ansiform lobule); Cr-II: crus II (ansiform lobule); CWM: cerebellar white matter; DN: dentate nucleus; EN: emboliform nucleus; Fl: Flocculus; FN: fastigial nucleus; GN: globose nucleus; MCP: middle cerebellar peduncle; Par-a: paramedian lobule, anterior part; Par-p: paramedian lobule, posterior part; PFl: paraflocculus; PIN: posterior interposed nucleus; SCP: superior cerebellar peduncle; Sim-a: simple lobule, anterior part; Sim-p: simple lobule, posterior part. For the abbreviations of the fissures (red text) separating the cerebellar lobules in the human (**C**), see Figure 7. Scale bars: 10 mm, applies to all scale bars in A-B; 50 mm, applies to C.

**Figure 7.**
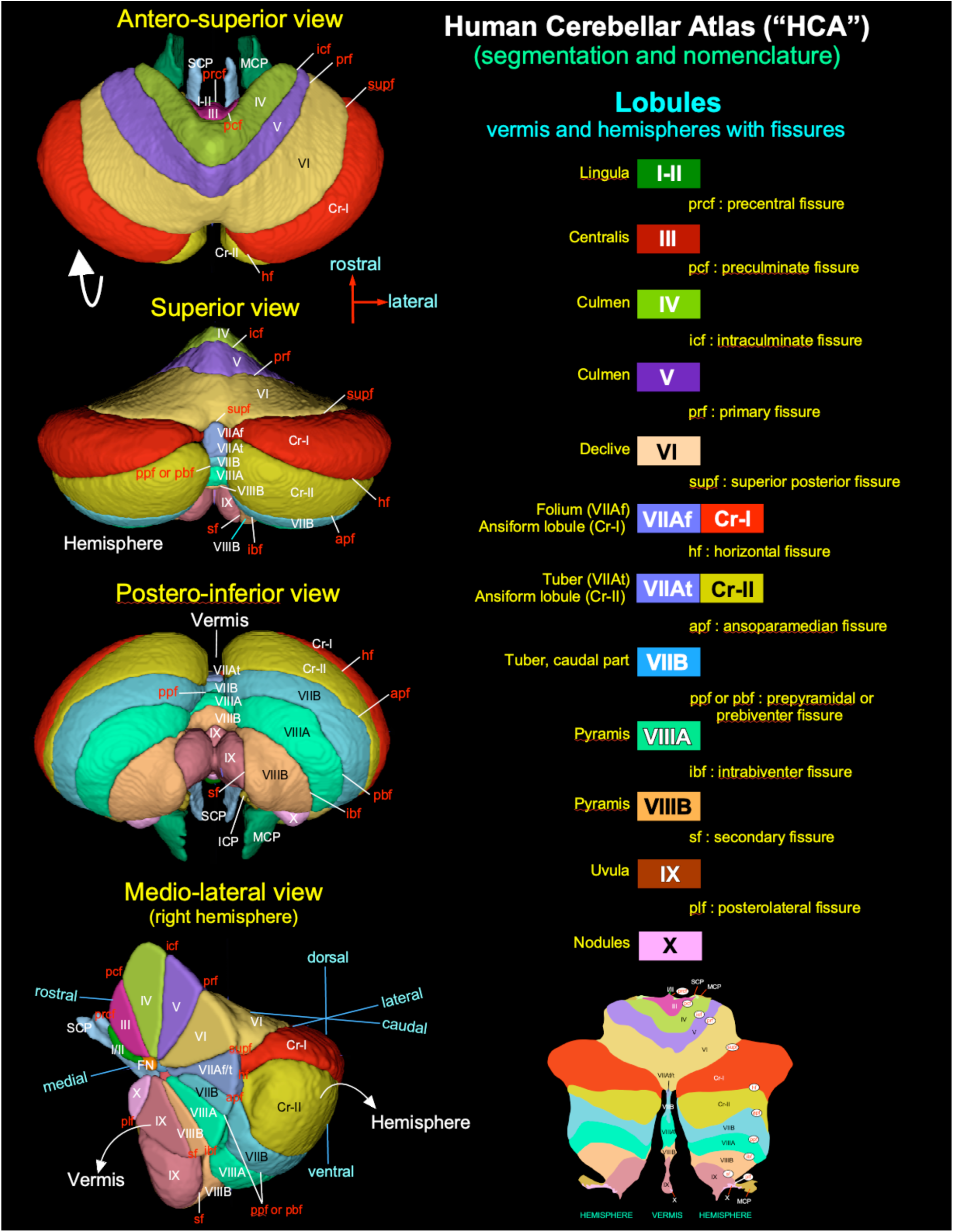
Human Cerebellar Atlas (HCA). The HCA provides a detailed three-dimensional asymmetric segmentation of the human cerebellum, including vermis and left and right hemispheric lobules, visualized from multiple angles. It incorporates detailed fissural and lobular definitions following Schmahmann et al. (*25*) and includes a flat map representation with asymmetric parcellation derived from HCA. The flat map (SUIT) cerebellar template is based on (*26*). Together, these resources enable precise characterization of human cerebellar lobular boundaries while preserving species-specific morphology, and support systematic cross-species comparisons across primates (Fig. 6A-C).

Across species, vermal lobules I-X were conserved and served as homologous anatomical anchors. In contrast, hemispheric territories within the posterior lobe (lobules VI-IX) exhibited progressive differentiation (Table 1). In the marmoset, the posterior hemispheric cortex was partitioned into discrete Simplex (Sim-a, Sim-p), Paramedian (Par-a, Par-p), and Copula of pyramis (Cop) lobules (Fig. 6A) (*27*). These territories correspond topologically to subdivisions of the human posterior cerebellar lobe: Sim regions align with hemispheric expansions of lobules VI–VII, whereas Par/Cop regions extend across territories corresponding to lobules VII-IX (Table 1; Figs. 6C, 7). Crus I and Crus II are present but remain relatively compact and lie in close spatial proximity to vermal VIIA, consistent with a less laterally elaborated hemispheric organization (Fig. 6A).

**Table 1.** Cross-species correspondence of posterior cerebellar lobules (VI-IX) reveals progressive hemispheric reorganization across primates. Correspondence between vermal and hemispheric cerebellar lobules in marmoset, macaque, and human. Vermal lobules (I-X) are aligned according to conserved anatomical nomenclature. In marmoset and macaque, hemispheric territories designated as Simplex (Sim-a, Sim-p), Paramedian (Par-a, Par-p), and the copula of pyramis (Cop) correspond topologically to subdivisions of the human posterior lobe (lobules VI-IX). Sim-a and Sim-p align with hemispheric expansions of lobule VI and anterior VII (VIIA), Par-a and Par-p occupy VII-VIII, whereas Cop occupy progressively more caudal hemispheric territories corresponding to human VIII-IX. Crus I and Crus II represent laterally expanded hemispheric domains associated with lobule VII (Figs. 6, 7). In macaque, these territories are spatially segregated from vermal VIIA and do not form a continuous surface extension, whereas in marmoset, partial spatial proximity is observed, reflecting reduced hemispheric differentiation. In humans, Crus I and Crus II constitute markedly expanded hemispheric territories with clear separation from the vermal cortex. Thus, alignment reflects developmental and topological homology within the posterior lobe rather than strict one-to-one lobular continuity, highlighting progressive lateral displacement and elaboration of posterior cerebellar hemispheres across primate evolution. Hemispheric labels reflect conventional usage in the respective atlases and literature and are intended to facilitate comparison rather than assert strict anatomical or evolutionary homology. Differences in hemispheric parcellation, including simple (Sim), crus (Cr), paramedian (Par), floccular (Fl), and parafloccular (PFl) regions, underscore divergence in cerebellar organization across primates. Anatomical nomenclature follows previously published studies (*23, 25, 27, 28*).

| Lobule (Vermis) | Hemisphere | Hemisphere | Hemisphere |
| --- | --- | --- | --- |
|  | **MARMOSET** | **MACAQUE** | **HUMAN** |
| I | – | – | – |
| II | II | II | II |
| III | III | III | III |
| IV | IV | IV | IV |
| V | V | V | V |
| VI | Simplex (Sim-a) | Simplex (Sim-a/p) | VI |
| VII (or VIIA) | Simplex (Sim-p), Crus I, Crus II | Simplex (Sim-p), Crus I, Crus II | Crus I |
| VII (or VIIB) | Paramedian (Par-a) | Paramedian (Par-p) | VIIB, Crus II |
| VIII (or VIIIA) | Paramedian (Par-a), Cop | Paramedian (Par-p), Cop | VIIIA |
| VIII (or VIIIB) | Paramedian (Par-p), Cop | Paramedian (Par-p), Cop | VIIIB |
| IX | Copula of pyramis (Cop) | Copula of pyramis (Cop) | IX |
| X (Nodulus) | Flocculus (Fl), Paraflocculus (PFl) | Flocculus (Fl), Paraflocculus (PFl) | Flocculus (or X) |

**Table 2.** Lobule-specific cerebellar volumes in marmoset, macaque, and human. Absolute volumes (mm³) are shown for individual cerebellar lobules and hemispheric subdivisions derived from species-specific atlases (Fig. 6). Lobules I-X follow conventional numbering. Crus I and Crus II (ansiform lobule) are listed separately. In marmoset and macaque, additional hemispheric subdivisions (simplex, paramedian, copula of pyramis, flocculus, and paraflocculus) are reported according to nonhuman primate anatomical conventions (*27, 28*). In humans, consistent with Schmahmann and colleagues (*25*), the flocculonodular lobe comprises lobule X (nodulus) and the flocculus; in our human cerebellar atlas (HCA), both the nodulus (vermal component) and the flocculus (hemispheric component) are incorporated within lobule X, and a distinct paraflocculus is not separately parcellated. Total cerebellar volume reflects the summed cortical subdivisions; deep cerebellar nuclei (DCN) volumes are shown separately.

| Volume of the cerebellum in the Marmoset, Macaque, and Human in mm<sup>3</sup> |  |  |  |  |  |  |  |  |  |  |  |  |  |  |  |  |  |  |  |  |  |
| --- | --- | --- | --- | --- | --- | --- | --- | --- | --- | --- | --- | --- | --- | --- | --- | --- | --- | --- | --- | --- | --- |
| Lobules | I | II | III | IV | V | VI | VII | VIII | IX | X | Crus I | Crus II | Sim_a | Sim_p | Par_a | Par_p | Cop | Fl | PFL | Total | DCN |
| **Marmoset** | 1.94 | 17.6 | 31.1 | 35.7 | 80 | 51.5 | 42.6 | 26.4 | 31.1 | 16.6 | 9.3 | 45.6 | 18 | 29.1 | 27.5 | 25.6 | 41.1 | 8.2 | 60.2 | **599.14** | **14.1** |
| **Macaque** | 32.9 | 135.1 | 390.2 | 416.1 | 847.7 | 285.3 | 214.7 | 180.1 | 219.9 | 76.4 | 616.1 | 486.3 | 282.9 | 395.9 | 378.4 | 436.4 | 735.2 | 77 | 478.8 | **6685.42** | **196.3** |
| **Human** | 161 |  | 1903.8 | 7791.6 | 9798 | 22330 | 15246 | 25230.6 | 9299.5 | 1310 | 38310.4 | 22780.2 |  |  |  |  |  |  |  | **154161.1** | **1384** |

In the macaque, hemispheric differentiation is more pronounced. The Simplex, Paramedian, and copula of pyramis lobules are clearly delineated and occupy expanded posterior hemispheric territories (Fig. 6B) (*28*). Crus I and Crus II form laterally positioned hemispheric domains associated with lobule VII but are spatially segregated from vermal VIIA and do not constitute a continuous surface extension (Fig. 6B). Thus, although developmentally related to lobule VII, Crus territories are anatomically displaced laterally in three-dimensional space. Humans exhibit the most extensive hemispheric reorganization: posterior cortex corresponding to lobules VI-IX is consolidated into continuous and markedly expanded lateral territories (Figs. 6C, 7). Crus I and Crus II form prominent hemispheric expansions clearly separated from vermal cortex, and posterior lobules VIIIA-IX are represented predominantly within the hemispheres, whereas anterior lobules (I-V) retain conserved vermal-hemispheric correspondence.

Together, these observations define a graded reorganization of posterior cerebellar cortex across primates, characterized by progressive lateral displacement and consolidation of lobules VI-IX into expanded hemispheric territories. Posterior regions, therefore, do not scale uniformly but become increasingly segregated from the vermis. Consistent with this structural redistribution, volumetric analyses demonstrate disproportionate expansion of posterior hemispheric subdivisions, particularly Crus I and Crus II, relative to anterior lobules and the deep cerebellar nuclei. We next quantified lobular volume allocation to determine how hemispheric differentiation scales across species and contributes to overall cerebellar expansion.

### Posterior cerebellar lobules and Crus I/II dominate cerebellar volume in humans

To determine whether the observed hemispheric reorganization corresponds to measurable shifts in volume, we quantified lobular scaling across species using atlas-derived measurements. Cerebellar lobular volumes were measured in marmoset, macaque, and human and grouped into major anatomical subdivisions to enable cross-species comparison (Fig. 8A: built-in table, B). Although the ansiform lobule (Crus I/II) is treated as a distinct subdivision, it occupies the lateral posterior hemisphere and is functionally associated with posterior cerebellar territories.

**Figure 8.**
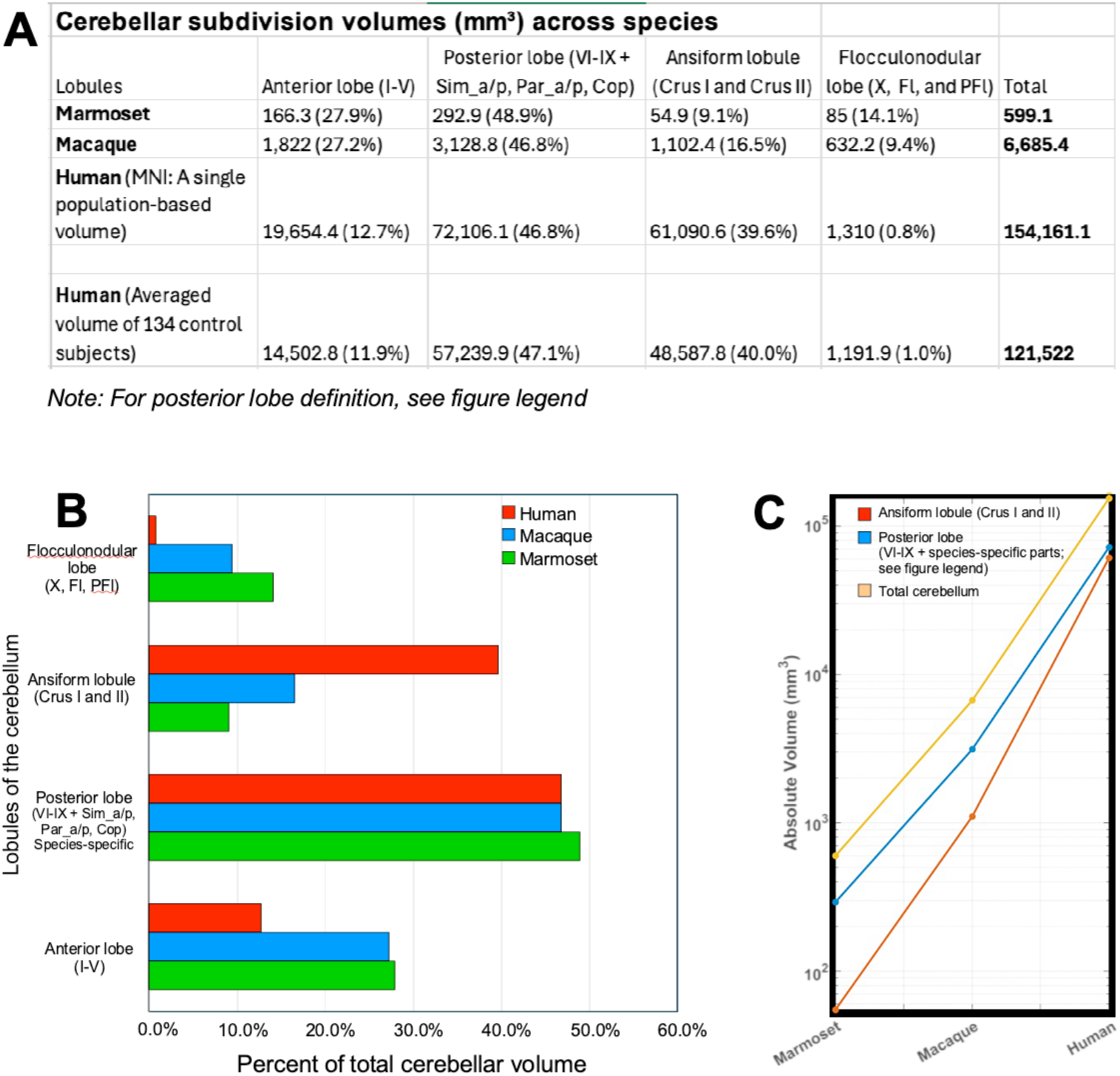
Disproportionate expansion of posterior cerebellar territories and the ansiform lobule (Crus I/II) across primates. **(A) Comparative cerebellar subdivision volumes across species.** Absolute volumes (mm³) and relative proportions of total cerebellar volume (%) are shown for major cerebellar subdivisions in marmoset, macaque, and human. Subdivisions include the anterior lobe (lobules I–V), posterior lobe, ansiform lobule (Crus I and Crus II), and flocculonodular lobe. In marmosets and macaques, the posterior lobe comprises lobules VI–IX together with the simplex (Sim_a/p), paramedian (Par_a/p), and copula of pyramis (Cop), which are anatomically defined as posterior lobe components in these species. In humans, the posterior lobe includes lobules VI–IX only and does not include Sim_a/p, Par_a/p, or Cop (see Table 1), as these structures are not parcellated as separate lobules in the Schmahmann cerebellar atlas (*25*). The flocculonodular lobe was defined according to species-specific anatomical conventions. In marmosets and macaques, it comprises lobule X (nodulus), the flocculus (FL), and the paraflocculus (PFL). In humans, consistent with Schmahmann et al. (*25*), the flocculonodular lobe includes lobule X (nodulus) and the flocculus; a distinct paraflocculus is not separately parcellated. In our human cerebellar atlas (HCA), both the nodulus (vermal component) and flocculus (hemispheric component) are incorporated within lobule X. The ansiform lobule (Crus I and Crus II) is presented as a separate subdivision to highlight its pronounced hemispheric expansion. Percent values indicate the proportion of total cerebellar volume for each species, and totals represent the summed volumes of all subdivisions. **(B) Relative cerebellar volume allocation across species.** The posterior lobe constitutes the largest subdivision in all species. However, the ansiform lobule (Crus I/II) shows a marked increase in proportional representation in humans, comprising ∼40% of total cerebellar volume, compared with substantially smaller proportions in the marmoset and macaque. This redistribution reflects preferential expansion of lateral posterior cerebellar territories. **(C) Absolute cerebellar volumes reveal differential scaling across species.** Log-scaled volumes demonstrate pronounced non-uniform expansion of cerebellar subdivisions. While total cerebellar volume increases substantially across species, Crus I/II exhibit the steepest scaling trajectory, expanding disproportionately relative to both total cerebellar volume and other subdivisions. Posterior lobe definitions differ across species (see panel A legend and main text), and values should be interpreted in the context of these anatomical distinctions.

In marmosets, the posterior lobe, defined as lobules VI–IX together with Sim_a, Sim_p, Par_a, Par_p, and Cop, was the largest subdivision, measuring 292.9 mm³ and accounting for 48.9% of total cerebellar volume, whereas the ansiform lobule (Crus I and Crus II) contributed 54.9 mm³ (9.1%). The anterior lobe (lobules I-V) measured 166.3 mm³ (27.9%), and the flocculonodular lobe measured 85 mm³ (14.1%). In macaques, the same posterior lobe definition yielded a volume of 3,128.8 mm³ (46.8%), while Crus I/II increased to 1,102.4 mm³ (16.5%). The anterior lobe accounted for 1,822 mm³ (27.2%) and the flocculonodular lobe for 632.2 mm³ (9.4%).

In humans, where the posterior lobe is defined as lobules VI-IX only, posterior lobe volume reached 72,106.1 mm³, comprising 46.8% of total cerebellar volume. Notably, the ansiform lobule expanded disproportionately to 61,090.6 mm³, accounting for 39.6% of the cerebellum. In contrast, the anterior lobe accounted for 12.7% and the flocculonodular lobe for only 0.8% of total cerebellar volume. Together, these results demonstrate a marked redistribution of cerebellar volume toward posterior cerebellar territories and, most prominently, the ansiform lobule (Crus I/II), indicating their dominance relative to other cerebellar subdivisions (Fig. 8B).

### Absolute (log-scale) expansion of posterior cerebellar territories

To distinguish proportional redistribution from absolute expansion across species, we examined cerebellar volumes on a logarithmic scale (Fig. 8C). Analysis of absolute volumes revealed pronounced differential scaling of posterior cerebellar territories. Because posterior lobe definitions differ across species, encompassing lobules VI-IX together with Sim_a/p, Par_a/p, and Cop in marmosets and macaques, but lobules VI-IX alone in humans, the reported values reflect species-specific anatomical organization.

The combined volume of these posterior regions increased from 292.9 mm³ in marmosets to 3,128.8 mm³ in macaques (∼10.7-fold increase) and further expanded to 72,106.1 mm³ in humans, corresponding to a ∼23-fold increase relative to macaques and a ∼246-fold increase relative to marmosets.

Crus I/II exhibited an even steeper scaling trajectory, increasing from 54.9 mm³ in marmosets to 1,102.4 mm³ in macaques (∼20.1-fold) and to 61,090.6 mm³ in humans, representing a ∼55-fold increase relative to macaques and a >1,100-fold increase relative to marmosets. By comparison, total cerebellar volume increased ∼11.2-fold from marmoset to macaque and ∼23.1-fold from macaque to human. Thus, on a logarithmic scale, Crus I/II emerge as the most rapidly expanding cerebellar subdivisions, exhibiting order-of-magnitude growth that exceeds both overall cerebellar scaling and the expansion of other posterior territories (Fig. 8C).

Together, these relative and absolute scaling patterns indicate selective, disproportionate expansion of posterior cerebellar territories, particularly the ansiform lobule (Crus I/II), which is preferentially linked to association cortices. This expansion provides a structural substrate for evolutionary reorganization of cerebro-cerebellar networks underlying human cognitive specialization.

### Relative Scaling of Cerebellar Lobules Is Preserved Across Native and Template MRI

To obtain population-level estimates of cerebellar lobular organization, we registered the Human Cerebellar Atlas (HCA), derived from a population-based MNI template (Fig. 6C) to in vivo T1-weighted MRI scans from 134 control subjects spanning a range of ages and both sexes (Table S1). This approach enabled consistent parcellation across individuals and the computation of average lobular volumes in native space. Aggregation of these measurements into major anatomical compartments revealed a proportional organization closely matching that of the MNI template (Fig. 8A). Notably, the posterior cerebellar lobules (lobules VI–IX, excluding Crus I/II) and Crus I and II together dominate the total cerebellar volume, accounting for a combined 87.1% of the structure. Specifically, the anterior lobe (lobules I–V) represented 11.9% of total volume, the posterior lobules comprised 47.1%, Crus I and II contributed 40.0%, and the flocculonodular lobe (lobule X) represented 1.0%. These values closely parallel those derived from the MNI template (12.7%, 46.8%, 39.6%, and 0.8%, respectively), indicating that the relative scaling of cerebellar compartments is preserved between a cohort average and a population-based reference.

Absolute cerebellar volumes derived from MNI space were consistently larger than those from the cohort average (Fig. 8A), in line with prior reports that template-based spatial normalization can introduce biases in volumetric estimates (*29, 30*). This effect likely arises from methodological factors such as nonlinear normalization, interpolation, and the use of an averaged anatomical template, which can systematically inflate regional volumes compared with cohort-specific measurements. Despite these differences in absolute scale, the close correspondence in proportional values suggests that relative compartmental organization is preserved across datasets. Together, these findings support the robustness of atlas-based parcellation and indicate that cerebellar scaling relationships remain largely consistent across individuals, demographic variation, and reference spaces.

## Discussion

### Cerebellar Input–Output Architecture Scales Nonuniformly Across Primates

The extent to which cerebellar input and output compartments scale in concert across primates remains unresolved. Using multimodal, high-resolution cross-species 3D atlases integrating iron-sensitive MRI with histology, we show that cerebellar cortical territories expand disproportionately relative to the deep cerebellar nuclei, revealing differential scaling of input-output architecture across marmoset, macaque, and human. Posterior hemispheric regions, particularly lobules VI-IX and Crus I/II, undergo preferential elaboration, whereas nuclear subdivisions exhibit selective rather than uniform enlargement. Together, these findings identify a systems-level organizational principle in which cerebellar cortex and nuclear output pathways scale nonuniformly, reshaping integration within distributed motor and association networks.

### Species-dependent Iron Patterns Shape DCN MRI Contrast

Species differences in DCN MRI contrast correspond closely with tissue iron content. T2 hypointensity in macaque and human DCN, relative to hyperintensity in marmosets, matched Perls’ iron staining and prior work linking MR signal to iron accumulation (*20, 31*). These findings indicate that iron contributes substantially to DCN contrast and support MRI as a biologically informed, noninvasive marker of regional iron content. Variation in iron-related contrast across species may reflect differences in metabolic demand within cerebellar output nuclei. Given iron’s central role in oxidative metabolism and mitochondrial function (*32, 33*), stronger labeling in macaque and human DCN is consistent with elevated bioenergetic requirements. Because iron strongly influences T2-weighted and susceptibility-based measures, these differences have important translational implications for interpreting DCN microstructure and pathology. Future work integrating quantitative susceptibility mapping could further refine links between MRI signal and cellular iron content (*34*).

### Histology-informed MRI Reveals Conserved and Species-specific DCN Organization

High-resolution, histology-validated MRI delineated the dentate (DN), interposed, and fastigial nuclei in marmosets, macaques, and humans (Figs. 2, 3). In nonhuman primates, T2-weighted imaging, supplemented with MTR and MAP-MRI metrics, reliably distinguished nuclei, while histological markers (SMI-32, NeuN, AChE, Nissl) confirmed boundaries and highlighted microanatomical features such as cell-sparse zones and differential neuropil labeling. Marmosets exhibited a small accessory fastigial nucleus (aFN), illustrating subtle species-specific variation. Extending this approach to humans, BigBrain-informed segmentation enabled clear identification of all major DCN, including the highly folded dentate nucleus, in postmortem datasets (*35*). Together, these results demonstrate that histology-guided MRI provides a robust framework for bridging nonhuman primate and human cerebellar mapping, enabling precise cross-species comparisons of structure, connectivity, and function (*3, 36, 37*).

### Dentate Nucleus Enlargement Supports Cerebellar Output Diversity

Quantitative volumetric analyses reveal selective enlargement of the dentate nucleus (DN) across species, with humans showing the most pronounced dominance. In marmosets, DCN volumes are relatively balanced, whereas macaques and humans exhibit disproportionate DN expansion, reflecting the DN’s central role as the principal output of the lateral cerebellum and its extensive connectivity with higher-order cortical systems (*3, 36, 38*). Our marmoset findings differ from those reported by Zhu and colleagues (*39*), who described the DN as the largest DCN subregion, exceeding the combined volume of the interposed and fastigial nuclei. In their study, the DN was delineated together with the anterior interposed (emboliform) nucleus, likely contributing to larger estimated DN volumes and DN-to-other-nuclei ratios. By contrast, our high-resolution MRI guided by histological ground truth enabled precise delineation of individual DCN subregions (Fig. 2), indicating that pronounced DN enlargement is not a defining feature of the marmoset but emerges progressively in larger-brained primates.

In humans, DN specialization is most pronounced, accounting for 1032.7 mm³ (86% of total DCN volume), whereas the fastigial and interposed nuclei contribute only minor proportions (Fig. 4). This pattern aligns with postmortem volumetric studies reporting DN contributions of ∼77% (*40*) and histology-based cytoarchitectonic analyses demonstrating a predominance of the DN over other nuclei (*14*). High-resolution imaging studies further corroborate this DN dominance (*41, 42*), although absolute DN volumes vary due to methodological differences, including partial-volume effects and limited resolution (*43*). Despite these differences, the relative dominance of the DN emerges as a robust feature of human cerebellar organization. Transcriptomic analyses indicate selective amplification of excitatory projection neurons within the DN, providing a cellular mechanism supporting increased output capacity (*44, 45*).

Beyond volumetric dominance, dentate nucleus enlargement has direct implications for cerebellar circuit architecture. As the principal output of the lateral cerebellum, the DN participates in closed-loop circuits linking cerebellar processing with distributed association cortices, including prefrontal and posterior parietal regions, through thalamic relays (*6, 38, 46–48*). Its internal organization into motor and nonmotor territories, featuring differentiated projection patterns to higher-order cortical regions (*38*), suggests that selective DN expansion increases the capacity and diversity of cerebellar output, supporting predictive and integrative computations extending beyond motor control (*3, 49*).

In contrast, the interposed and fastigial nuclei, which are primarily associated with sensorimotor and axial functions (*50–52*), with the exception of the posterior interposed nucleus, scale more modestly across species. This divergence reinforces the view that cerebellar evolution preferentially amplifies lateral hemispheric-dentate circuits interfacing with association cortical networks rather than uniformly expanding all output channels.

### Progressive Posterior Hemispheric Specialization Shapes Cerebellar Integration

Complementing selective dentate enlargement, posterior cerebellar hemispheric territories (lobules VI-IX and Crus I/II) undergo disproportionate expansion relative to anterior and vermal lobules. In marmosets, posterior cortex is partitioned into discrete Simplex, Paramedian, and Copula of pyramis lobules, with Crus I/II relatively compact and partially associated with vermal VIIA. In macaques, Crus territories become laterally segregated from vermal cortex, forming distinct hemispheric domains. In humans, posterior lobules consolidate into expansive lateral territories, with Crus I/II emerging as dominant hemispheric structures largely dissociated from vermal counterparts.

Quantitative analyses show that posterior hemispheric components account for an increasingly large fraction of total cerebellar volume from marmoset to macaque to human, whereas anterior lobules and vermal territories scale more modestly. This preferential expansion parallels reciprocal cerebro-cerebellar interactions involving association cortices, mediated through pontine input pathways and thalamic output channels (*5, 36, 53*), providing structural substrates for higher-order integrative processing. Functional imaging in humans shows that these posterior territories, particularly Crus I/II, are engaged in executive, attentional, and social-cognitive tasks (*12*). While deep nuclei scale selectively, the disproportionate cortical expansion supports the interpretation that cerebellar elaboration primarily enhances integrative processing capacity rather than uniformly increasing output channels.

### Behavioral and Evolutionary Implications of Selective Cerebellar Remodeling

Comparative analyses indicate that cerebellar cortical and nuclear compartments scale nonuniformly, reflecting distinct functional specializations across primates. Medial vermal and interposed-fastigial circuits contribute to posture, locomotion, and gaze control (*50, 52*), supporting arboreal navigation in nonhuman primates and refined postural and visuomotor regulation in humans. In contrast, posterior hemispheric territories (lobules VI-IX and Crus I/II) exhibit disproportionate expansion, accompanied by scaling of the dentate nucleus. These lateral hemispheric-dentate circuits are extensively integrated with distributed association networks across the cerebral cortex (*3, 36*), consistent with broader evidence for preferential expansion of cortico-cerebellar systems across anthropoid primates (*9, 54, 55*). Although these anatomical patterns do not imply a strict causal link between structure and behavior, they support the view that cerebellar organization is selectively biased toward circuits interfacing with association networks.

Evolutionarily, this bias reflects mosaic, non-isometric remodeling of cortico-cerebellar systems rather than uniform scaling with overall brain size. Phylogenetic evidence indicates coordinated expansion of lateral cerebellar hemispheres with association cortex (*9, 55*), and these circuits are implicated in predictive processing, sequencing, and adaptive control beyond motor function (*3, 36*). Ecological pressures, including arboreal locomotion and increasing social complexity, likely contributed to this trajectory by favoring enhanced visuomotor integration and flexible behavioral planning (*54*). Notably, the disproportionate expansion of posterior cerebellar cortex relative to deep nuclei points to evolutionary pressures that preferentially enhance input-side processing capacity while maintaining more constrained output channels. Together, these findings advance a model in which cerebellar evolution is tightly coupled to the expansion of distributed association networks, with the most pronounced elaboration in humans.

### Limitations of the study

Despite the novel insights provided by our cross-species cerebellar atlases, several limitations should be acknowledged. First, the study is based on a modest number of specimens per species (n = 5), which may limit the ability to fully capture inter-individual variability, including potential sex- or age-related differences. Nevertheless, the combination of high-resolution MRI and histological delineation for each case ensures detailed and reliable anatomical measurements, supporting the consistency of the observed cross-species patterns. Second, subtle differences in tissue processing, scan resolution, or segmentation protocols across species could introduce minor measurement variability. Third, our focus on volumetric and lobular comparisons does not directly assess microstructural, connectivity, or functional differences, which are essential for understanding the full scope of cortico-cerebellar integration. Finally, although the study highlights broad patterns of posterior expansion and altered cortical-nuclear scaling, evolutionary interpretations remain correlational; additional developmental, functional, and behavioral data will be needed to link structural changes to cognitive or sensorimotor adaptations. These limitations underscore the need for future studies integrating larger sample sizes, multimodal functional imaging, and finer-scale histological analyses to refine our understanding of cerebellar evolution in primates.

## Conclusion

Using multimodal MRI-histology atlases spanning marmoset, macaque, and human, we show that cerebellar expansion is characterized by nonuniform scaling of input-output architecture rather than homogeneous enlargement. Posterior cortical territories expand disproportionately relative to anterior lobules and deep cerebellar nuclei, accompanied by progressive lateral consolidation of hemispheric domains. Subdivisions associated with lobule VII, particularly Crus I and II, undergo marked hemispheric elaboration, whereas anterior vermal-hemispheric correspondence remains comparatively conserved. Importantly, cortical enlargement is not matched by proportional expansion of the deep cerebellar nuclei, revealing a shift in the balance between cortical input and nuclear output. These multimodal atlases provide a powerful resource for comparative neuroanatomy, enabling precise correspondence between imaging features and underlying cellular architecture across species. Evolutionarily, cerebellar enlargement in primates reflects selective architectural remodeling rather than uniform growth. Preferential expansion of posterior hemispheric cortex and dentate output pathways mirrors the broader elaboration of higher-order cortical systems, revealing coordinated reorganization of cerebro-cerebellar circuits. Rather than scaling passively, the cerebellum emerges as an actively restructured hub, supporting integrative networks underlying complex human cognition.

## Materials and Methods

### Non-human primate Specimens and Ethical Compliance

Brains from two adult common marmosets (*Callithrix jacchus*, male 340 g; female 283 g) and one adult rhesus macaque (*Macaca mulatta*, male 13.55 kg) were obtained from the National Institute of Mental Health (NIMH/NIH) for our MRI and histological analyses. These specimens were already perfusion-fixed as part of prior studies at NIMH; we did not perform any live experiments or perfusions on these animals for our research. The marmosets had previously been used in transgenic studies, and the macaque in behavioral studies. Animals had been anesthetized with sodium pentobarbital and perfused transcardially with 0.5 L heparinized saline, followed by 1-2 L (marmosets) or 4 L (macaque) of 4% paraformaldehyde in 0.1 M phosphate buffer (pH 7.4). Brains were removed, photographed, post-fixed for 8-24 h, and stored in PBS with sodium azide until MRI. Specimens were rinsed in PBS before scanning to reduce residual fixative effects. All procedures conformed to the Guide for the Care and Use of Laboratory Animals (National Research Council) and were approved by the NIMH/NIH Institutional Animal Care and Use Committee.

### Ex Vivo MRI Acquisition in Marmosets

For MRI acquisition, fixed brains (cases 1-2) were positioned in 3D-printed molds (Fig. 9) within 30-mm cylindrical containers filled with Fomblin and vacuum-degassed to eliminate air bubbles. Samples were sealed and scanned on a Bruker 7T/300 mm horizontal-bore MRI system using a 30-mm quadrature Millipede coil (ExtendMR). The optimal MRI acquisition parameters differed slightly between the two cases. Detailed descriptions of marmoset and macaque sample preparation and MRI protocols have been reported previously (*19, 20*).

**Fig. 9.**
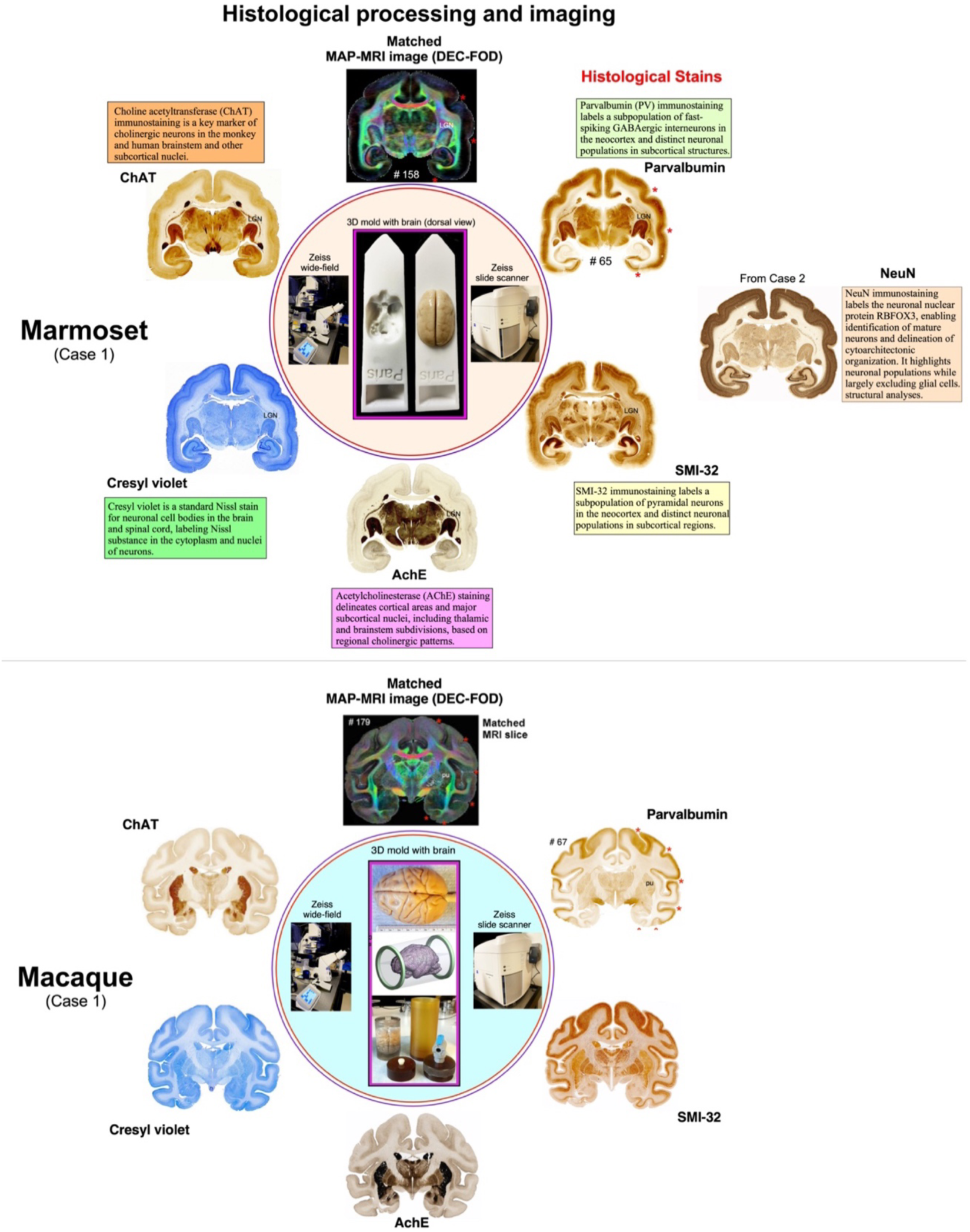
Complementary histological staining and MRI correspondence in the marmoset and macaque. In marmoset Case 1, serial coronal frozen sections (50 μm) spanning the rostrocaudal extent of the cerebral hemispheres, from frontal to occipital pole, were collected and assigned to interleaved staining series to preserve rostrocaudal correspondence across complementary histological stains. The illustrated sections shown at the level of the anterior temporal cortex include Nissl staining, acetylcholinesterase (AChE) histochemistry, and immunohistochemistry for parvalbumin (PV), SMI-32 (nonphosphorylated neurofilament H), and choline acetyltransferase (ChAT). In the marmoset (Case 2) and the macaque, additional NeuN immunostaining and Perls’ Prussian blue histochemistry for iron detection were performed (iron stain not shown). High-resolution brightfield images of stained sections were acquired using a Zeiss wide-field microscope and a Zeiss Axioscan Z1 slide scanner (inset). Histological sections were manually matched to corresponding MAP-MRI images (top), and additional structural MRI contrasts from the same specimen, using conserved anatomical landmarks, including sulcal fundi, gyral crowns, ventricular contours, and deep gray nuclei. Note the correspondence of sulci (red stars), gyri, and deep brain structures in both MRI and histology sections (“Red stars”). This cross-modal correspondence enabled landmark-guided delineation of subcortical territories, including the deep cerebellar nuclei, within a common three-dimensional reference framework. The inset also shows the specimen-specific 3D brain mold used to preserve ex vivo brain geometry during MRI acquisition.

For cases 1 and 2, ex vivo MAP-MRI data were acquired at 150-µm isotropic resolution using a 3D diffusion spin-echo EPI sequence. Case 1 was acquired with a 288 × 184 × 184 imaging matrix; FOV, 4.32 × 2.76 × 2.76 cm; TE/TR = 48/650 ms; 10 segments; 1.33 partial Fourier acceleration; δ/Δ = 6/28 ms; and 256 DWIs across 11 b-value shells (100-10,000 s/mm²) with 7-40 uniformly distributed gradient directions per shell. Case 2 was acquired with a 256 × 160 × 160 imaging matrix; FOV, 3.84 × 2.40 × 2.40 cm; TE/TR = 43/1400 ms; 8 segments; 1.25 partial Fourier; δ/Δ = 8/20 ms; and 112 DWIs across six b-value shells (100-10,000 s/mm²) with 3-36 directions per shell; two averages were acquired at b = 10,000 s/mm² to improve SNR. In both cases, diffusion directions were uniformly sampled on the unit sphere (*56, 57*).

Magnetization transfer (MT)- prepared 3D gradient echo images were also acquired at 150-µm isotropic resolution (TE/TR = 3.7/37 ms; 15° flip angle) using a 2-kHz offset, 12.5-ms Gaussian saturation pulse (6.74 µT peak amplitude; 540° flip angle), with four averages for MT-on and MT-off scans. Total acquisition times were 85 h 20 min (case 1) and 74 h 40 min (case 2) for MAP-MRI, and 11 h 8 min (case 1) and 8 h 31 min (case 2) for MT imaging.

A third perfusion-fixed marmoset brain sample (Case 3, female, 289 g) was prepared for MRI in a manner similar to the other two specimens. The sample was scanned on a vertical-bore 7T Bruker MRI system using a 30 mm millipede RF coil and a 3D diffusion spin-echo EPI sequence. Data were acquired with 86 µm isotropic resolution.

### Ex Vivo MRI Acquisition in the Macaque

A 3D structural MRI of the fixed ex vivo macaque brain was first obtained and reoriented to the stereotaxic plane of the D99 atlas (*19, 58*). The aligned volume was used to 3D-print a custom cylindrical brain mold for stable positioning during scanning. The specimen was placed in a 70-mm cylindrical container filled with Fomblin, vacuum-degassed for 4 h to remove air bubbles, sealed, and scanned on a Bruker 7T/300 mm horizontal-bore MRI system using a 72-mm quadrature RF coil. All MRI volumes were registered to the D99 atlas using affine and nonlinear transformations.

MAP-MRI data were acquired at 200-µm isotropic resolution (375 × 320 × 230 matrix; FOV, 7.5 × 6.4 × 4.6 cm; TE/TR = 50/650 ms; 8 segments; 1.33 partial Fourier acceleration; δ/Δ = 6/28 ms) with 112 DWIs across six b-value shells (100-10,000 s/mm²) and 3-36 uniformly distributed directions per shell. MT-prepared 3D gradient echo images were acquired at 250-µm isotropic resolution (TE/TR = 3.5/37 ms; 15° flip angle) using a 2-kHz offset, 12.5-ms Gaussian saturation pulse (6.74 µT peak amplitude; 540° flip angle), with two averages for MT-on and MT-off scans. Total acquisition times were 93 h 20 min (MAP-MRI) and 6 h 18 min (MT).

### Diffusion modeling and microstructural metrics in marmoset and macaque

Post-processing of MRI data was identical across species. Diffusion-weighted images were processed using the TORTOISE software package (*59*) to correct for Gibbs ringing, signal drift, and distortions arising from magnetic field inhomogeneity and eddy currents, with the MTR image serving as the structural reference. The mean apparent propagator was estimated voxelwise using a fourth-order MAP series approximation implemented in MATLAB. From the diffusion tensor model, we derived fractional anisotropy (FA), mean diffusivity (MD), axial diffusivity (AD), radial diffusivity (RD), and tensor shape coefficients (CL, CP, CS). We derived MAP-MRI metrics including propagator anisotropy (PA), return-to-origin (RTOP), return-to-axis (RTAP), and return-to-plane (RTPP) probabilities, non-Gaussianity (NG), and the non-diffusion-weighted amplitude image (T2-weighted contrast). Orientation distribution functions (ODFs) and fiber ODFs were estimated and visualized using MRtrix3 (*60*). MTR maps were computed from MT-on and MT-off images and provided high gray-white matter contrast for reliable registration to diffusion volumes.

### Histology and immunohistochemistry

Following MRI acquisition, brains were cryoprotected, sectioned, and processed for histological staining. Histological procedures were identical for marmoset and macaque specimens and are described in detail elsewhere (*19, 20*) (Fig. 9). Briefly, serial sections were processed for Nissl staining, acetylcholinesterase (AChE) histochemistry, and immunohistochemistry using antibodies against parvalbumin (PV), SMI-32 (nonphosphorylated neurofilament H), and choline acetyltransferase (ChAT). In the marmoset case 2 and the macaque, an additional staining for NeuN and Perls’ Prussian blue (iron) was performed. Immunoreactions were visualized using the avidin–biotin complex method with diaminobenzidine as the chromogen. Sections were mounted, dehydrated, cleared, and coverslipped using standard procedures. Detailed protocols and antibody information are described previously (*19, 20*), and the software used in this study is summarized in Table 3.

**Table 3.**
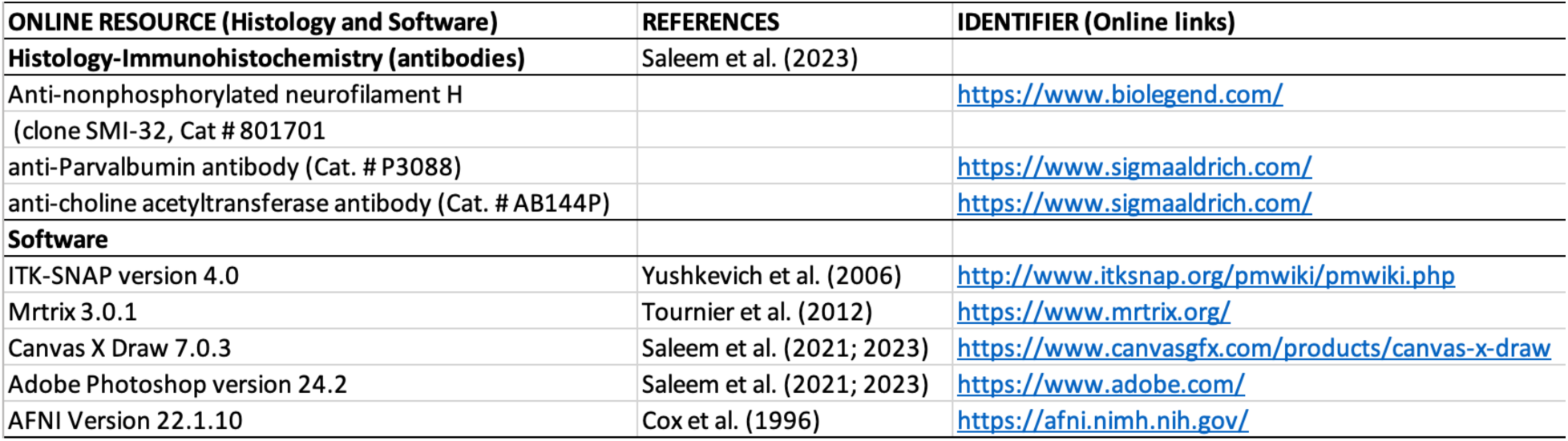
Antibodies and Computational Tools for MRI-Histology Integration, Segmentation, and Atlas Construction. This table lists the primary antibodies used for immunohistochemical staining, as well as the software tools and in-house pipelines used for image processing, cerebellar region segmentation, 3D visualization, and atlas reconstruction. (e.g., ITK-SNAP for manual segmentation; and AFNI with SUMA for surface visualization and spatial normalization).

| ONLINE RESOURCE (Histology and Software) | REFERENCES | IDENTIFIER (Online links) |
| --- | --- | --- |
| **Histology-Immunohistochemistry (antibodies)** | Saleem et al. (2023) |  |
| Anti-nonphosphorylated neurofilament H<br>(clone SMI-32, Cat # 801701) |  | https://www.biolegend.com/ |
| anti-Parvalbumin antibody (Cat. # P3088) |  | https://www.sigmaaldrich.com/ |
| anti-choline acetyltransferase antibody (Cat. # AB144P) |  | https://www.sigmaaldrich.com/ |
| **Software** |  |  |
| ITK-SNAP version 4.0 | Yushkevich et al. (2006) | http://www.itksnap.org/pmwiki/pmwiki.php |
| Mrtrix 3.0.1 | Tournier et al. (2012) | https://www.mrtrix.org/ |
| Canvas X Draw 7.0.3 | Saleem et al. (2021; 2023) | https://www.canvasgfx.com/products/canvas-x-draw |
| Adobe Photoshop version 24.2 | Saleem et al. (2021; 2023) | https://www.adobe.com/ |
| AFNI Version 22.1.10 | Cox et al. (1996) | https://afni.nimh.nih.gov/ |

### Data analysis

High-resolution images of all stained sections were acquired using a Zeiss wide-field microscope and an Axioscan Z1 slide scanner (5× objective) (Fig. 9). Digital images were adjusted for brightness and contrast and manually aligned to the corresponding diffusion-derived maps (DTI and MAP-MRI metrics), estimated T2-weighted (non-diffusion-weighted) images, and MTR volumes to facilitate visualization and 3D segmentation (see below) of cerebellar lobules, deep cerebellar nuclei, and the cerebellar peduncles (Figs. 2, 3, 6). Accurate alignment was achieved without resampling the MRI volumes. Precise orientation of the intact brain specimen relative to sagittal MRI images was performed prior to sectioning, enabling consistent matching of sulci, gyri, and deep ROIs across MRI and histological sections (Fig. 9) (*19, 20*).

### Cerebellar segmentation, 3D atlas generation, and volumetric analysis

In marmoset and macaque, deep cerebellar nuclei (dentate nucleus [DN], anterior interposed nucleus [AIN], posterior interposed nucleus [PIN], and fastigial nucleus [FN]) and vermal and hemispheric subdivisions of cerebellar lobules were manually segmented from ex vivo coronal MRI volumes (primarily T2-weighted and complementary MAP-MRI contrasts) using ITK-SNAP (*61*). MRI-defined boundaries were validated against matched high-resolution histological sections (SMI-32, NeuN, AChE, and Nissl stains) and informed by previously described cerebellar anatomical frameworks in these species, with nomenclature aligned to prior studies (*23, 27, 28*). Segmented regions were reconstructed in three dimensions and visualized using ITK-SNAP (*61*) and SUMA (*62*) to define spatial relationships among cerebellar cortical and nuclear subdivisions. These reconstructions formed the basis of the Marmoset Cerebellar Atlas (MCA) and Rhesus Macaque Cerebellar Atlas (RMCA). Ex vivo MCA and RMCA segmentations were subsequently transformed into species-specific stereotaxic space and aligned to population-based in vivo templates for atlas generation (Fig. 6A, B). All segmentations were manually reviewed and refined to ensure accurate areal extent and architectonic boundaries of cerebellar lobules and deep nuclei, with particular reference to fissural landmarks and histological features. Spatial normalization and segmentation were performed in AFNI (*62, 63*) using pipelines optimized for high-resolution ex vivo imaging. The resulting population-based in vivo templates constitute the standard MCA and RMCA (Fig. 6A, B) used for quantitative analyses and will be made publicly available through the AFNI platform (*link forthcoming*).

For humans, cerebellar lobules and deep cerebellar nuclei (DN; emboliform nucleus [EN, corresponding to AIN]; globose nucleus [GN, corresponding to PIN]; and FN) were segmented with reference to established anatomical descriptions (*14, 25*). The T1-weighted MNI_icbm152 nonlinear template (*64, 65*) served as the stereotaxic reference space. Because deep nuclei are not well resolved in standard T1-weighted MNI templates, high-resolution cytoarchitectonic data from the BigBrain dataset (*35*) were co-registered to MNI space to refine nuclear boundaries (Figs. 2C, F; figs. S3, S5). In addition, in vivo diffusion data, including direction-encoded color (DEC) volumes from the Human Connectome Project (*66*), were nonlinearly registered to this space (fig. S5B) to aid delineation of cerebellar peduncles and intracerebellar fiber orientations (analyses will be reported in a separate manuscript). Integration of these multimodal datasets enabled the generation of the Human Cerebellar Atlas (HCA) (Fig. 6C).

To generate a flatmap representation of the human cerebellum, we first aligned the MNI 2009c cerebellar volume to the SUIT cerebellar template (*26*) using nonlinear warping implemented in *3dQwarp* (AFNI). The resulting deformation field was then applied to the HCA (Human Cerebellar Atlas) to bring it into SUIT space. Next, the transformed atlas was projected onto the SUIT surface representation using the *vol2surf* plugin, sampling voxel values along the cortical ribbon between the white matter and pial surfaces with a modal (most frequent value) mapping approach. Finally, the surface-projected atlas was smoothed on the pial surface using 10 mm modal smoothing to reduce local discontinuities while preserving discrete parcellation boundaries.

Each atlas includes detailed segmentation of vermal and hemispheric lobules and deep cerebellar nuclei (DCN), standardized according to Paxinos/Larsell/Schmahmann nomenclature (*23–25, 27, 28*), and provides multiplanar visualizations in coronal, axial, and sagittal orientations in stereotaxic or MNI coordinates (Fig. 6A-C). For the HCA, detailed fissural landmarks separating lobules are also provided (Fig. 7). Homologous lobules and nuclei were identified across marmoset, macaque, and human templates based on structural landmarks and histological validation, enabling reliable cross-species comparisons (Table 1). The 3D cerebellar atlas templates in these species, and the script for atlas registration of in vivo scans, are now available for download through the AFNI and SUMA websites at https://afni.nimh.nih.gov/…………………. (*link forthcoming*). The AFNI software can install this simply with the @Install_SAM_Marmoset command.

For consistency, volumetric measurements of cerebellar lobules and deep cerebellar nuclei were obtained from finalized 3D atlas segmentations derived from species-specific, population-based templates (Figs. 6, 7; Table 2; Figs. 4, 5, 8). This approach provides a standardized framework for cross-species comparison while minimizing biases arising from individual variability. The resulting digital atlases define stereotaxic reference volumes and surface parcellations suitable for comparative cerebellar mapping and integration into neuroimaging pipelines.

### Additional Cases for Cross-Species Quantitative DCN Analysis

To enhance cross-species volumetric comparisons of the deep cerebellar nuclei (DCN), we supplemented our primary datasets with independently acquired cases from marmosets, macaques, and humans, yielding five cases per species (Fig. 4, top panel). In the marmoset, beyond the three ex vivo T2-weighted MRI datasets generated in this study (Cases 1-3), we incorporated the population-averaged in vivo MBM_v3.0 T2-weighted template (200 μm resolution; (*67*) and a high-resolution ex vivo MBM_v5.0 dataset (80 μm resolution; (*39*) (fig. S4).

In the macaque (Fig. 4, top panel), we combined our ex vivo T2-weighted dataset (200 μm resolution; Case 1) with an ex vivo T2-weighted (G12) dataset registered to the NMT v2.0 population-based template (*68*), and three quantitative susceptibility mapping (QSM) datasets (400 μm resolution) provided by Atsushi Yoshida (*34*). QSM contrast improved visualization and delineation of DCN subregions. In both marmosets and macaques, DCN boundaries were segmented serially on T2-weighted and QSM sections with direct reference to closely matched histological sections obtained from our own ex vivo cases (SMI-32, NeuN, and Nissl staining), ensuring species-specific anatomical fidelity.

For humans, in addition to the MNI T1-weighted template, we analyzed four independent T2-weighted MRI volumes (800 μm isotropic resolution) spanning different ages and both sexes (Fig. 4, top panel), obtained from the OpenNeuro (*69*). DCN segmentation was performed on serial T2-weighted sections using the BigBrain template registered to subject space as a histological reference (*35*) (fig. S4). Together, this multi-source, multimodal framework enabled robust evaluation of DCN volumetry across species while minimizing template- and cohort-specific bias.

## Acknowledgments

This work was supported by the Intramural Research Program of the *Eunice Kennedy Shriver* National Institute of Child Health and Human Development; the Intramural Research Program of the National Institute of Mental Health (NIMH); and the NIH BRAIN Initiative project “Connectome 1.0: Developing the next-generation human MRI scanner for bridging studies of the micro-, meso-, and macro-connectome” (1U01EB026996-01). We thank Drs. Betsy Murray and Richard Saunders, and Alex Cummins (Laboratory of Neuropsychology, NIMH), for providing perfusion-fixed rhesus monkey brain for our experiments, and to James Pickel and the Transgenic Core Facility at NIMH for providing the perfusion-fixed marmoset brains used in this study. We also thank the Human Brain Collection Core (HBCC), NIMH/NIH, for providing the fixed human brain specimen and the Neurophysiology Imaging Facility Core (NIMH, NINDS, NEI) for MRI scanning. We further acknowledge Vincent Schram and his team at the Microscope Imaging Core (MIC) at NICHD for assistance with high-resolution imaging of histological sections. Histological processing of brain tissue was performed by Dr. Du and colleagues at FD NeuroTechnologies (Columbia, MD, USA).

## Author contributions

Kadharbatcha S. Saleem: Corresponding author, designed the study, coordinated the project, prepared the specimen for MRI and histology; mapped, segmented, and verified all the anatomical regions of interest with reference to MRI and histology; generated a new cerebellar 3D atlas template/Table, made illustrations; and wrote, edited, and streamlined the manuscript (Conceptualization, Data curation, Formal analysis, Investigation, Methodology, Project administration, Software, Supervision, Validation, Visualization, Writing-original draft, Writing-review & editing).

Alexandru V. Avram: Conducted all MRI experiments, including MAP-MRI scans, processed and analyzed all MRI data, helped with atlas data registration, and wrote and edited the manuscript (Data curation, Methodology, Writing-review & editing).

Daniel Glen: Integrated the atlas dataset into the AFNI and SUMA software packages, wrote the code for atlas region regularization, helped with atlas data registration, and edited the manuscript (Methodology, Software, Visualization, Writing-review & editing).

Peter J. Basser: Helped design the study, edited the manuscript, and provided research resources (Conceptualization, Funding acquisition, Resources, Writing-review & editing).

---

**Fig. S1.**
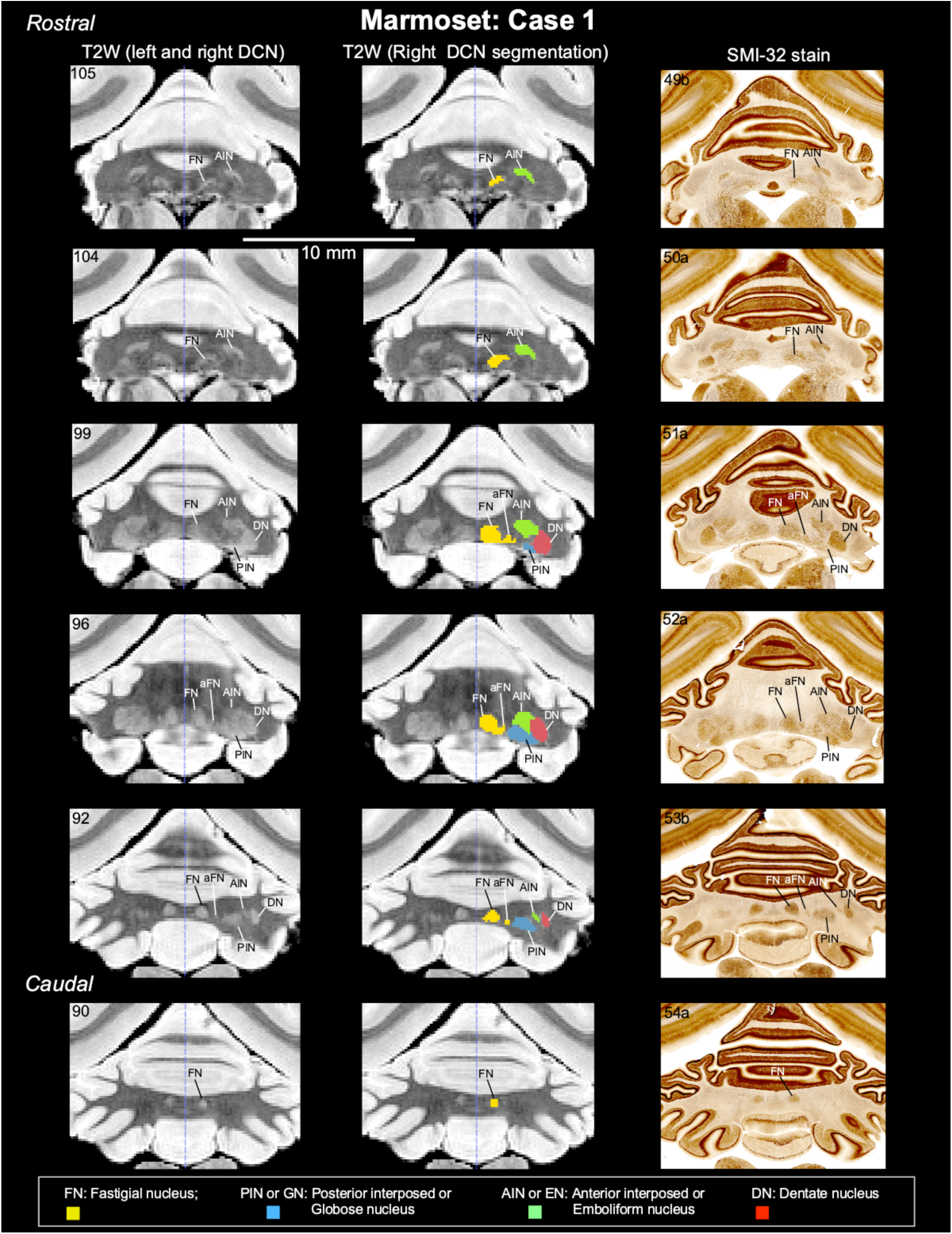
Histology-guided segmentation of the deep cerebellar nuclei (DCN) in marmoset, Case 1. Rostrocaudal T2-weighted MRI slices at 150 µm resolution show the left and right DCN with hyperintense signal (left column). Segmentation of DCN subregions on the right hemisphere is shown in the middle column. These subregions were delineated with reference to matched histology sections from the same case stained with SMI-32 (right column).

**Fig. S2.**
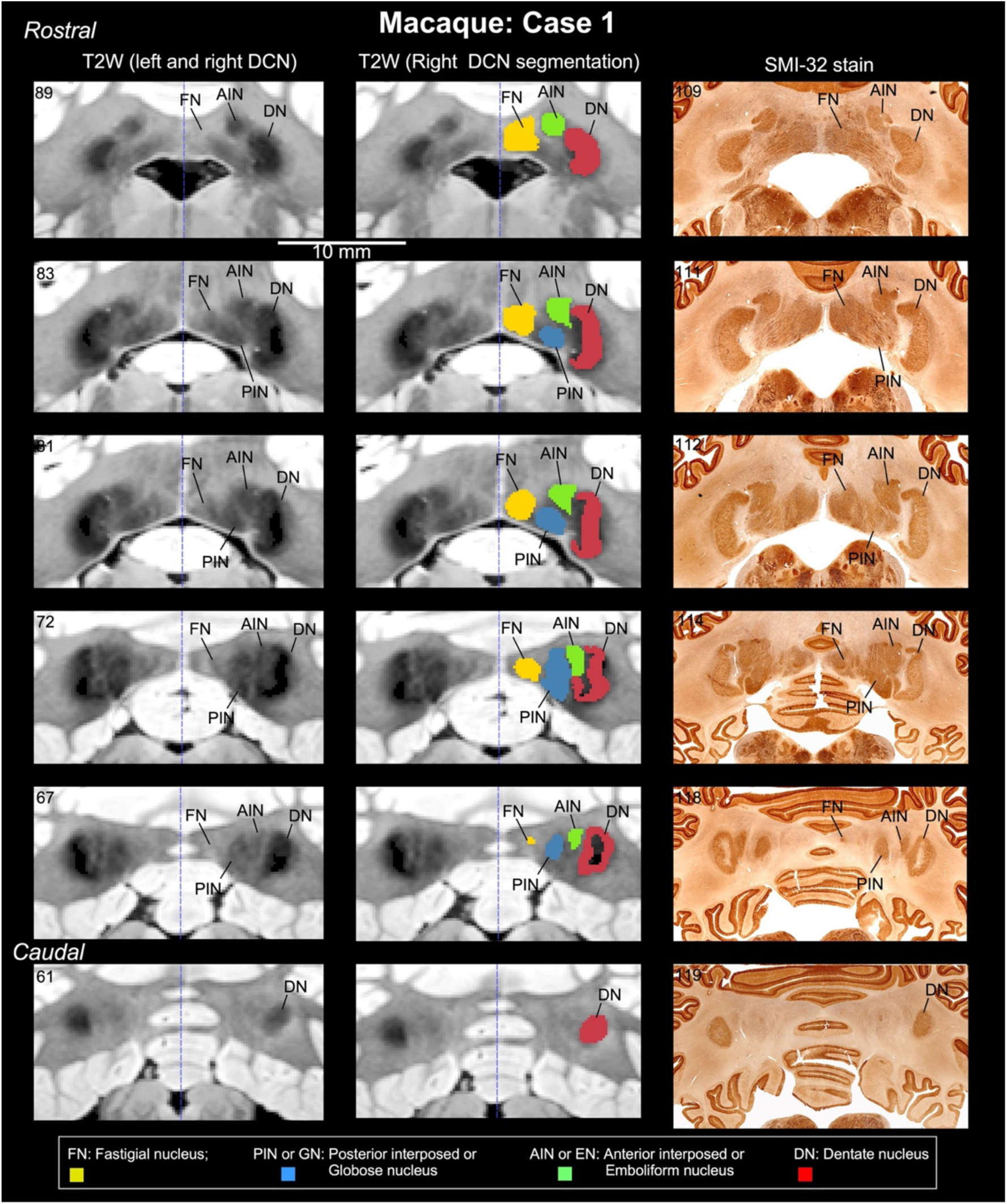
Histology-guided segmentation of the deep cerebellar nuclei (DCN) in macaque, Case 1. Rostrocaudal T2-weighted MRI slices at 200 µm resolution show the left and right DCN with hypointense signal (left column). Segmentation of DCN subregions on the right hemisphere is shown in the middle column. These subregions were delineated with reference to matched histology sections from the same case stained with SMI-32 (right column).

**Fig. S3.**
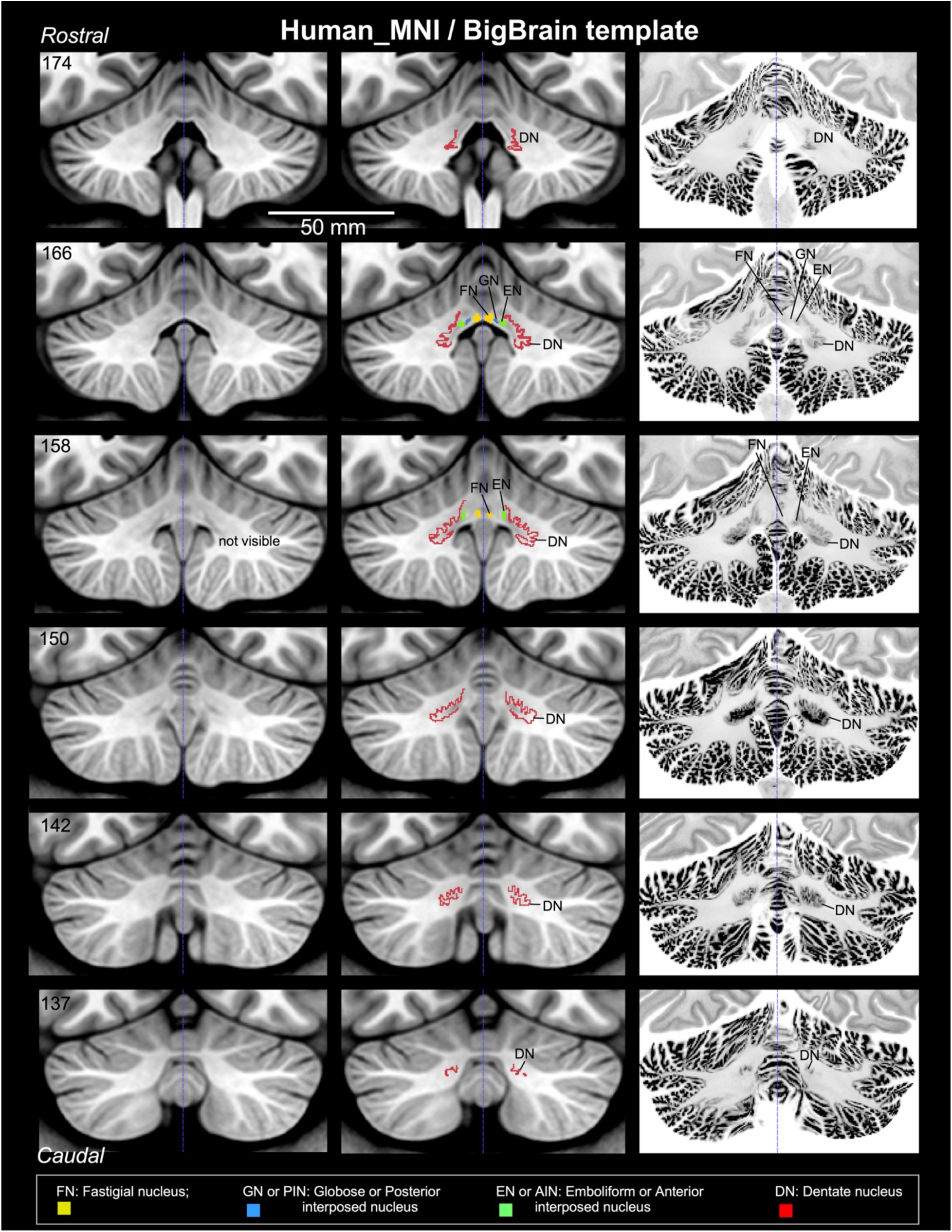
Histology-guided segmentation of the deep cerebellar nuclei (DCN) in humans. Rostrocaudal T1-weighted MRI slices from the ICBM-152 MNI template at 500 µm resolution do not provide sufficient contrast in the cerebellar white matter to delineate DCN subregions, unlike T2-weighted images in marmoset and macaque (left column). Segmentation of DCN subregions on the right hemisphere is shown in the middle column. These subregions were delineated with reference to matched histology sections stained with Nissl from the BigBrain template registered to MNI space (right column).

**Fig. S4.**
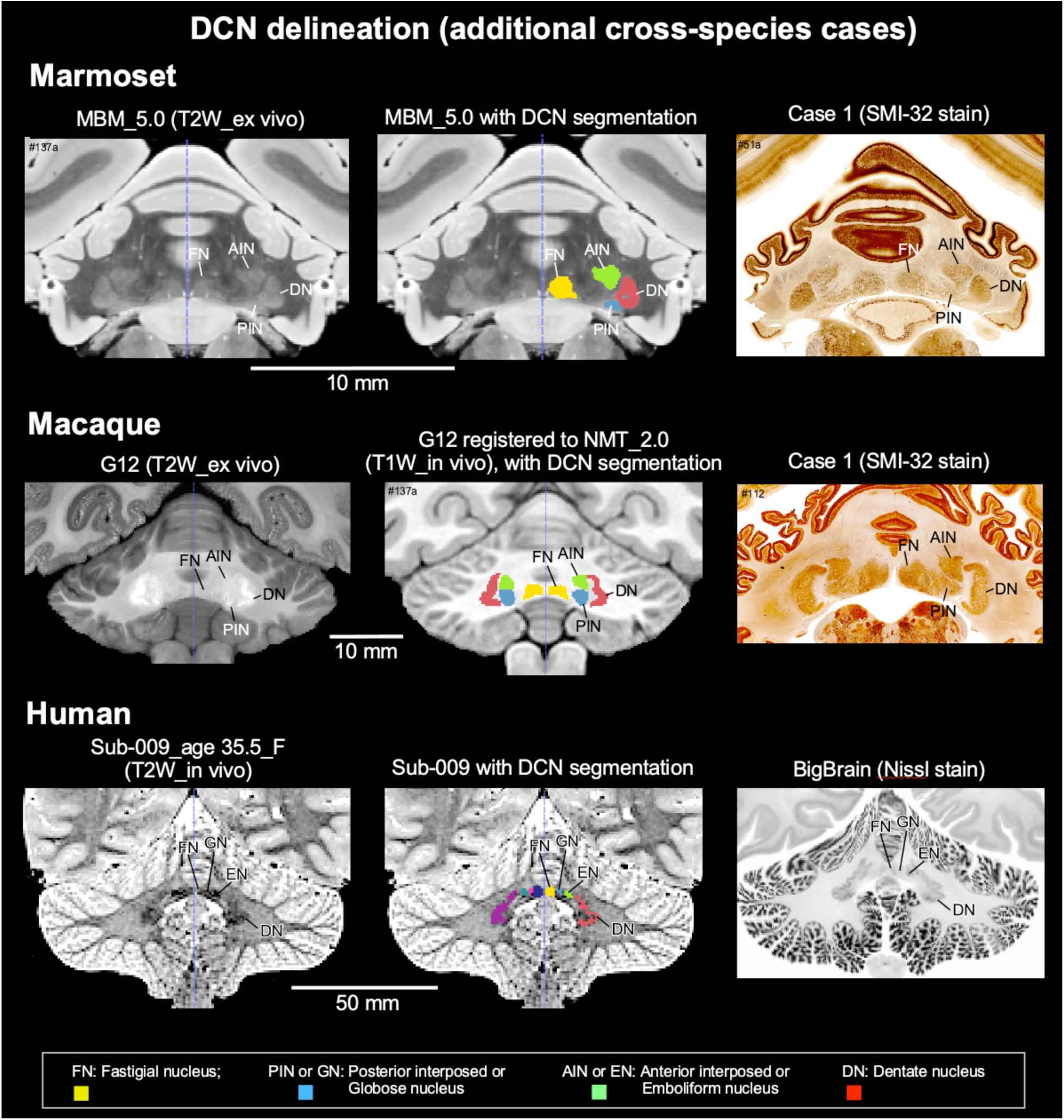
Additional cross-species cases for DCN segmentation and quantitative analysis. To strengthen cross-species volumetric comparisons of the deep cerebellar nuclei (DCN), we supplemented our primary datasets with independently acquired cases from marmosets, macaques, and humans, yielding five cases per species (Fig. 4, top panel; see Materials and Methods). **Top row:** High-resolution ex vivo MBM_v5.0 T2-weighted MRI (80 µm; Zhu et al., 2023) shows hyperintense DCN subregions, with segmentation on the right hemisphere. Borders were verified using a matched SMI-32–stained histology section from a different marmoset (Case 1). **Middle row:** Contrast-adjusted ex vivo T2-weighted macaque dataset (G12) was registered to the T1-weighted NMT v2.0 population-based template (Hartig et al., 2021), showing bilateral DCN segmentation. Boundaries were confirmed using a matched SMI-32 histology section from a different case, ensuring species-specific anatomical fidelity. **Bottom row:** One of the four human in vivo T2-weighted MRI volume (800 µm isotropic; OpenNeuro; Chai et al., 2025) (Fig. 4, top panel) shows hypointense DCN signal with bilateral segmentation, verified against a matched BigBrain histology slice. This multi-source, multimodal framework enables robust cross-species evaluation of DCN volumes while reducing template- and cohort-specific biases.

**Fig. S5.**
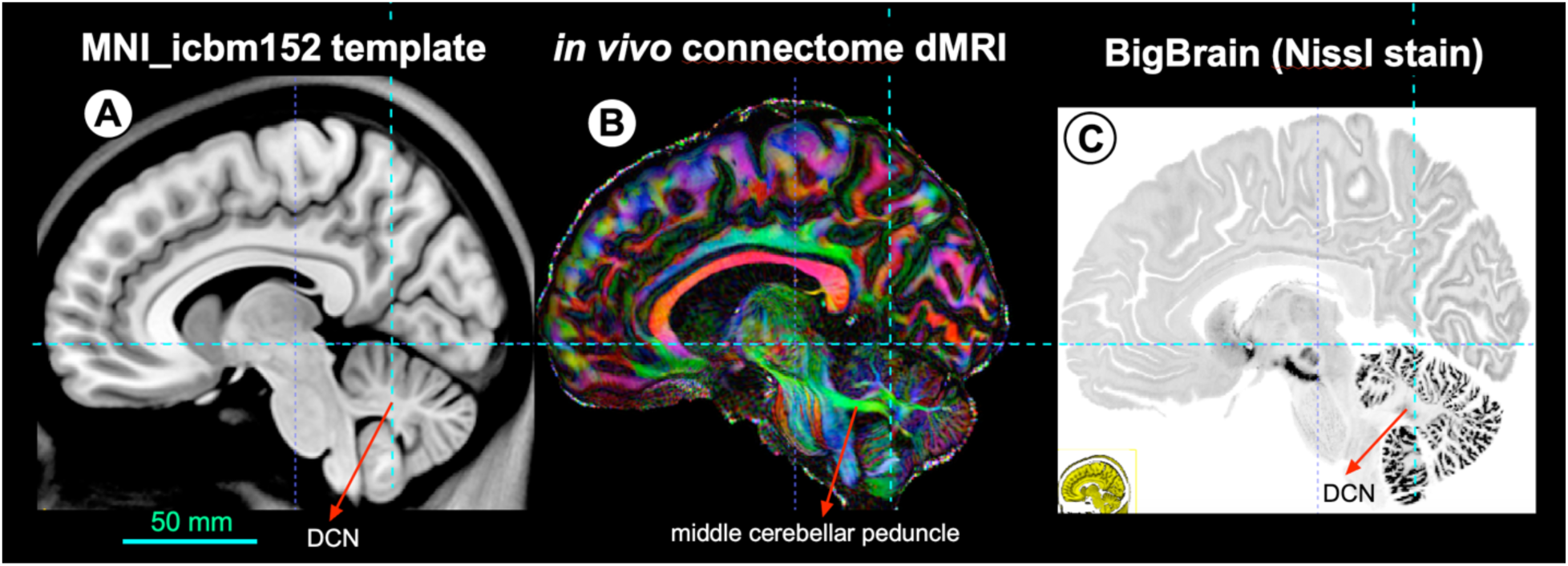
Multimodal integration for generation of the Human Cerebellar Atlas (HCA). Because the deep cerebellar nuclei (DCN) are not reliably delineated in standard T1-weighted MNI templates (A), high-resolution cytoarchitectonic data from the BigBrain dataset (C) were co-registered to MNI space to refine nuclear boundaries. In parallel, in vivo diffusion MRI data from the Human Connectome Project, including direction-encoded color (DEC) volumes (B), were nonlinearly registered to the same space to delineate cerebellar peduncles and intracerebellar fiber orientations. The integration of structural MRI, histology, and diffusion data enabled precise definition of DCN subregions and white matter architecture, forming the basis of the Human Cerebellar Atlas (HCA).

**Table S1.** Cerebellar lobular volumes in a normative cohort (n = 134). Mean gray matter volumes (mm³) for individual cerebellar lobules in the right and left hemispheres, as well as total lobular volumes (combined hemispheres), are reported for lobules I–X, including VIIA–f/t, VIIB, VIIIA, VIIIB, and Crus I/II, in 134 healthy participants aged 12–52 years (both sexes). Lobules are grouped anatomically: the anterior lobe (I–V), posterior lobe (VI–IX), and lobule X (flocculonodular lobe) reported separately. Interindividual variability is expressed as standard deviation. Posterior lobules (VI–IX) and Crus I/II exhibit the largest volumes, whereas anterior lobules (I–V) and lobule X are comparatively smaller (Fig. 8A). The cohort was compiled from multiple publicly available datasets, including Frontiers-QC, ABIDE1 (Kennedy Krieger), Trinity from ABIDE1, KUL3-ABIDE2, FCP Baltimore (Kennedy Krieger), and OpenNeuro (DS000220 and DS000243).

| Cerebellar lobules | Volume (Right cerebellum) | Volume (Left cerebellum) | Volume (total; Mean) | stdev |
| --- | --- | --- | --- | --- |
| I/II | 42.4903 | 68.356 | 110.8463 | 0.0971437 |
| III | 668.593 | 714.634 | 1383.227 | 0.0986591 |
| IV | 3113.95 | 2793.49 | 5907.44 | 0.117828 |
| V | 2676.21 | 4425.09 | 7101.3 | 0.0687519 |
| VI | 8756.92 | 7831.44 | 16588.36 | 0.0870774 |
| VIIA-f/t | 160.295 | 308.902 | 469.197 | 0.0605252 |
| VIIB | 6160.48 | 6037.32 | 12197.8 | 0.10656 |
| VIIIA | 5369.54 | 5878.55 | 11248.09 | 0.110855 |
| VIIIB | 4486.22 | 5004.08 | 9490.3 | 0.116704 |
| IX | 3336.7 | 3909.5 | 7246.2 | 0.0797186 |
| X | 558.121 | 633.77 | 1191.891 | 0.0917725 |
| Cr_I | 15549.5 | 15095.5 | 30645 | 0.0656142 |
| Cr_II | 8871.5 | 9071.29 | 17942.79 | 0.100844 |
| **Total number of subjects: 134** |  |  |  |  |
| **Age range: 12-52 (M/F)** |  |  |  |  |

